# Improved Photocleavable Proteins with Faster and More Efficient Dissociation

**DOI:** 10.1101/2020.12.10.419556

**Authors:** Xiaocen Lu, Yurong Wen, Shuce Zhang, Wei Zhang, Yilun Chen, Yi Shen, M. Joanne Lemieux, Robert E. Campbell

**Affiliations:** Department of Chemistry, University of Alberta, Edmonton, Alberta, T6G 2G2, Canada; Department of Biochemistry, Membrane Protein Disease Research Group, University of Alberta, Edmonton, Alberta, T6G 2H7, Canada; Talent Highland, The First Affiliated Hospital, Xi’an Jiaotong University, Xi’an, Shaanxi, 710061, China; Department of Chemistry, Stanford University, Stanford, California, 94305, United States; Department of Biochemistry, University of Alberta, Edmonton, Alberta, T6G 2H7, Canada; Department of Chemistry, The University of Tokyo, Tokyo, 113-0033, Japan

## Abstract

The photocleavable protein (PhoCl) is a green-to-red photoconvertible fluorescent protein that, when illuminated with violet light, undergoes main chain cleavage followed by spontaneous dissociation of the resulting fragments. The first generation PhoCl (PhoCl1) exhibited a relative slow rate of dissociation, potentially limiting its utilities for optogenetic control of cell physiology. In this work, we report the X-ray crystal structures of the PhoCl1 green state, red state, and cleaved empty barrel. Using structure-guided engineering and directed evolution, we have developed PhoCl2c with higher contrast ratio and PhoCl2f with faster dissociation. We characterized the performance of these new variants as purified proteins and expressed in cultured cells. Our results demonstrate that PhoCl2 variants exhibit faster and more efficient dissociation, which should enable improved optogenetic manipulations of protein localization and protein-protein interactions in living cells.

## Introduction

As a burgeoning range of biological techniques, optogenetics, which involves the use of light and genetically encodable light-sensitive proteins, enables unprecedented levels of control of numerous biological processes ranging from cellular activities to animal behaviours^1–3^. Genetically encodable light-sensitive proteins, as a key component of optogenetic actuators, are typically engineered from natural photoreceptors^4^. The current repertoire of engineered optogenetic actuators can be divided into the following categories^4,5^: (i) opsin-based light-activatable channels or pumps (e.g., ChR2 [Ref. 6], eNpHR^7^, eBR^8^ and OptoXR^9^); (ii) proteins with light-induced allosteric change (e.g., LINuS^10^, LEXY^11^ and PA-Rac1 [Ref. 12]); (iii) proteins with light-induced dimerization (e.g., TULIPs^13^, Magnets^14^, CRY2-CIB1 [Ref. 15], PhyB-PIF^16^, BphP1-PpsR2 [Ref. 17] and Dronpa 145N^18^); and (iv) photocleavable protein (PhoCl)^5^. As this summary reveals, PhoCl is the sole member of a distinct class of optogenetic actuators and uniquely enables irreversible optogenetic activation via a mechanism that requires cleavage of a covalent bond.

The first generation PhoCl was engineered from a circularly permuted (cp) green-to-red photoconvertible fluorescent protein (FP) that, when illuminated with ∼400 nm violet light, undergoes a main chain-breaking β-elimination at the green chromophore^19^. The bond cleavage produces a large N-terminal empty barrel fragment and a small C-terminal peptide fragment which spontaneously dissociate (as schematically represented in **Fig. 1a**,**b**). Due to this property, PhoCl1 can be used as a relatively simple yet versatile tool for covalent caging of a protein of interest that can be subsequently activated irreversibly by light. We previously demonstrated the use of PhoCl1 for the engineering a photoactivatable Cre recombinase, Gal4 transcription factor, and a protease that was used in turn to activate the ion channel Pannexin-1 (PanX1)^5^. Others have further expanded the applications of PhoCl1 to include photo-responsive biomaterials^20,21^, photo-control of cell-to-cell mechanotransduction^22^, and light-induced protein phase separation^23^. Although PhoCl1 has been applied in a growing number of cell physiology and biomaterials applications, a substantial drawback has been the relatively long half-time (*t*_1/2_) of fragment dissociation (*t*_1/2_ ∼ 500 s)^5^.

**Figure 1.**
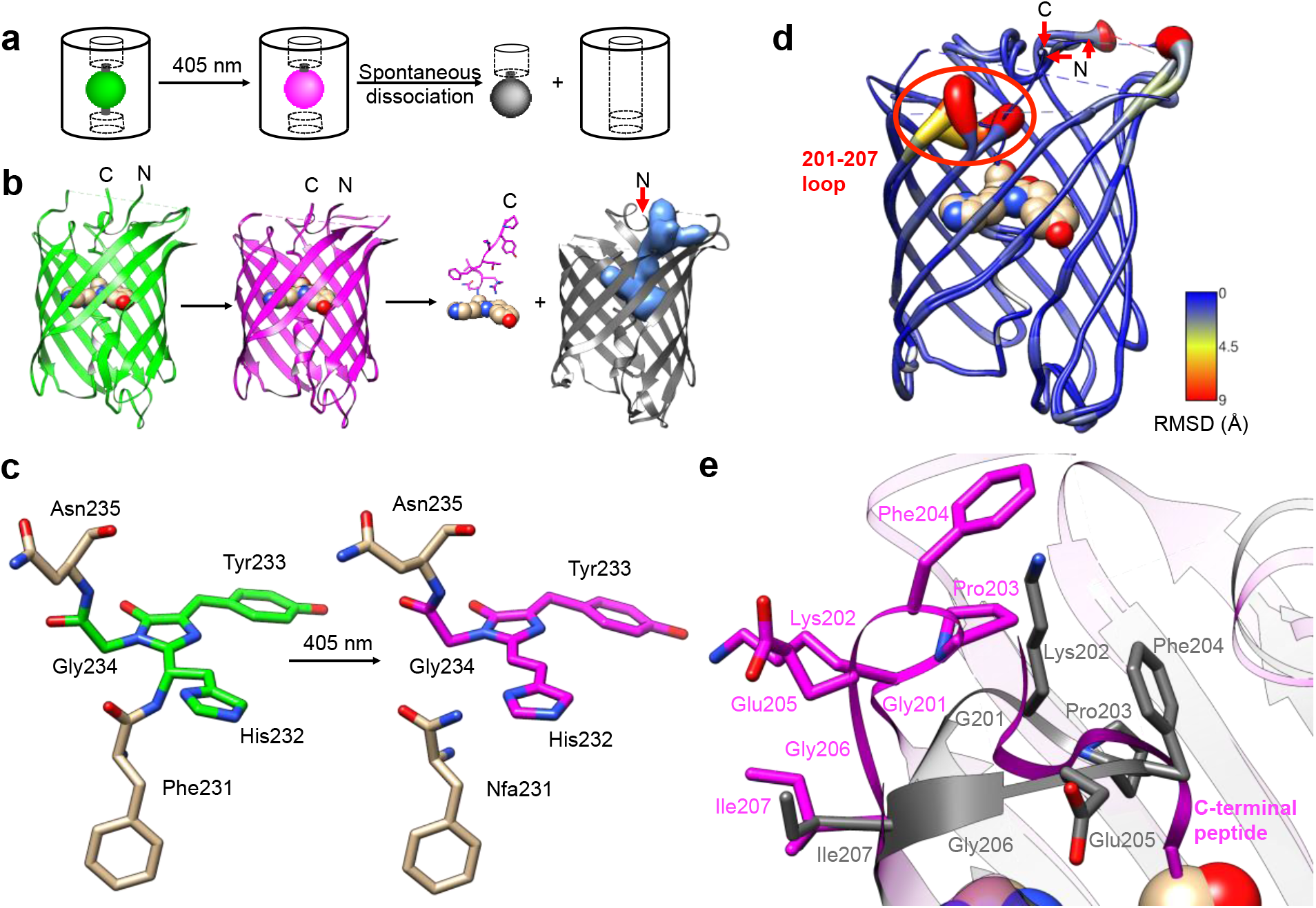
Overview of PhoCl1 structure and function. (**a**) Schematic of PhoCl photoconversion and dissociation. (**b**) Representation of PhoCl1 structures. Green state of PhoCl1: green structure (PDB ID: 7DMX), the red state: magenta structure (PDB ID: 7DNA), and the cleaved PhoCl1 empty barrel: grey structure (PDB ID: 7DNB). Water-filled cavity is represented as a blue space-filling volume. Dissociated peptide fragment is shown as sticks for the peptide portion and spheres for the chromophore portion. (**c**) The experimentally-determined structure of the PhoCl1 chromophore in the green and red states. (**d**) Structure alignment of the PhoCl1 red state and empty barrel colored by RMSD values. The RMSD values are indicated by scale bar and coil thickness (thicker coil indicates higher RMSD). (**e**) Zoomed-in representation of the 201-207 loop in the aligned structures of the red state (magenta) and empty barrel (grey).

To increase the general utility and applicability of PhoCl, we undertook the development of second generation PhoCl variants (PhoCl2) with improved rate and efficiency of dissociation. To achieve this goal, we first solved the X-ray crystal structures of the green state, the red state, and the cleaved empty barrel of PhoCl1. Then, using structure-guided engineering and directed evolution, in combination with a NanoLuc luciferase^24^-based screening assay^25^, we engineered the PhoCl2c variant with higher dissociation contrast ratio and the PhoCl2f variant with faster dissociation rate. Compared to the original PhoCl1, these PhoCl2 variants exhibit improved dissociation as purified proteins and in cell-based experiments. We further applied PhoCl2c for the control of light-induced cell apoptosis in living cells. These results establish the improved variants, PhoCl2c and PhoCl2f, as useful tools for optogenetic control of protein localization and function in live cells.

## Results

### Crystal structures of PhoCl1

To gain a better understanding of the structural changes associated with photoconversion and peptide dissociation, and to aid our protein engineering efforts, we determined the X-ray crystal structures of the PhoCl1 green state (2.1 Å resolution), red state (2.3 Å resolution), and the empty barrel fragment (2.8 Å resolution) (**Fig. 1b** and **Supplementary Table 1**). The structure of the peptide fragment was not determined experimentally and is schematically represented in **Fig. 1b** only for the sake of clarity.

To produce the PhoCl1 red state crystals, we first crystallized the purified PhoCl1 in its default green state, and then subjected these crystals to *in situ* partial photoconversion (**Supplementary Fig. 1a**). By illumination with 405 nm violet light (15 s light with LED array, 3.46 mW/mm^2^), green state PhoCl1 crystals changed from a visibly yellow color to a visibly orange-red color, indicating successful photoconversion. The visibly orange-red color of the photoconverted PhoCl1 appeared to be indefinitely stable as it was maintained for at least one month. We did not observe color loss in the context of the photoconverted red state crystal, suggesting no peptide dissociation occurred, presumably due to the very high concentration of protein and/or steric constraints imposed by crystal packing.

To purify the PhoCl empty barrel protein for crystallization, the intact PhoCl protein with a C-terminal 6× His purification tag was captured on a column of Ni-NTA agarose beads and then photoconverted on the column with repeated violet light illumination (405 nm LED array, 3.46 mW/mm^2^) to induce photocleavage and dissociation. The eluent containing the PhoCl1 empty barrel was collected, further purified by gel filtration chromatography, and then the protein was crystallized.

The green and red state structures revealed the changes in the chromophore structure associated with photoconversion. The green chromophore of PhoCl1 is generated from the autogenic post-translational modification of His232, Tyr233, and Gly234 (residues numbering are consistent with the numbering used in the X-ray crystal structure of the green state PhoCl1), which are structurally aligned with Ser65, Tyr66, and Gly67, respectively, of the canonical *Aequorea victoria* green FP (GFP; PDB ID: 1EMA)^26,27^. Determination of the protein structure using the partially photoconverted crystal revealed a mixture of green and red PhoCl1 states. As expected on the basis of the known photoconversion mechanism^19,28,29^, the red state of PhoCl1 was cleaved at the amide bond connecting Phe231 to His232 (**Fig. 1c**). Other than the changes in the vicinity of Phe231 and the chromophore, the overall structures of the green and red states are essentially identical.

To our surprise, the best fit of the electron density obtained from the partially photoconverted crystal was achieved by modeling only 2 of the 6 monomers in the asymmetric unit in the red state. Initial attempts to model all of the 6 monomers in the asymmetric unit as a mixture of green and red chromophore structures resulted in unsatisfactory fits to the experimental electron density. The best fit of the electron density was achieved by modeling 2 monomers as red state chromophores (chains A and B), and 4 monomers as green state chromophores (chains C, D, E, F). It remains unclear why 2 of 6 monomers were preferentially photoconverted upon illumination, rather than all 6 monomers experiencing equal photoconversion and each existing as a mixture of 33% red state and 67% green state. We speculate that efficient Förster resonance energy transfer (FRET) between the closely packed proteins in the crystal may have funneled energy into those monomers that were in the least favorable orientation to act as FRET donors and/or the most favorable orientation to act as FRET acceptors. These monomers may have been more likely to undergo photoconversion due to their higher probability of excitation or longer excited state lifetimes.

The structure of the PhoCl1 empty barrel revealed substantial conformational changes relative to the red state, as expected given the complete absence of the C-terminal fragment. Photocleavage and dissociation leaves a water-filled cavity where the peptide fragment previously resided in the interior of the empty barrel. To better visualize the conformational changes between PhoCl1 red state and the empty barrel, the root-mean-square deviation (RMSD) for all residues (that is, the average RMSD for all atoms in each residue) between the two structures was calculated, and the resulting RMSD values were used to color the cartoon coil alignment representation and adjust the coil diameter (thicker coil indicates higher RMSD) (**Fig. 1d**). This representation revealed that the most substantial, and observable, conformational changes occur in the 201-207 loop (highlighted in **Fig. 1d**). Specifically, in the empty barrel structure, the 201-207 loop region is “folded in” towards the center of the barrel, relative to its position in the red state structure (**Fig. 1e**). Due to this reorientation of the 201-207 loop into the barrel, the cavity in the empty barrel structure differs in shape from the volume that had been occupied by the C-terminal peptide (**Supplementary Fig. 1b,c**). We speculate that this conformational change helps to stabilize the dissociated empty barrel structure by partially filling the hydrophobic void generated by dissociation of the small peptide fragment. Accordingly, we reasoned that introducing mutations within or near the 201-207 loop region may affect the rate and efficiency of peptide dissociation, and we therefore focussed our efforts on this key region in our subsequent engineering.

### Molecular dynamics (MD) simulation of the dissociation process

To gain insight into the dissociation process of PhoCl1 at the molecular level, an adaptive steered molecular dynamics (ASMD) scheme (**Supplementary Fig. 2a**) was employed to investigate the dissociation pathway. Overall, 3 structures with distinct conformations of the modeled cp linker (i.e., the linker that connects the original C- and N-termini, **Supplementary Fig. 2b**) served as the initial coordinates of the ASMD, all of which were taken from a 210 ns unstrained canonical molecular dynamics (cMD) trajectory during which no spontaneous dissociation occurred (**Supplementary Fig. 2a,b** and **Supplementary Movie 1-4**). During the ASMD, the distance between the centres-of-mass of the N-terminal barrel and the C-terminal peptide were increased by 1 Å over 1 ns for each of 40 iterative stages, to give a total final displacement of 40 Å. Each stage was repeated 100 times from which an averaged potential mean of force was calculated and the replicate closest to the average was taken as the starting point for the next stage. We simulated the dissociation through 40 such stages without manually specifying the route or direction of dissociation (**Supplementary Fig. 2**). In all 3 distinct ASMD replications, the C-terminal peptide dissociated through the space between the flexible cp linker (residue 147-172) and the 201-207 loop (**Fig. 2a,b** and **Supplementary Movies 2-4**).

**Figure 2.**
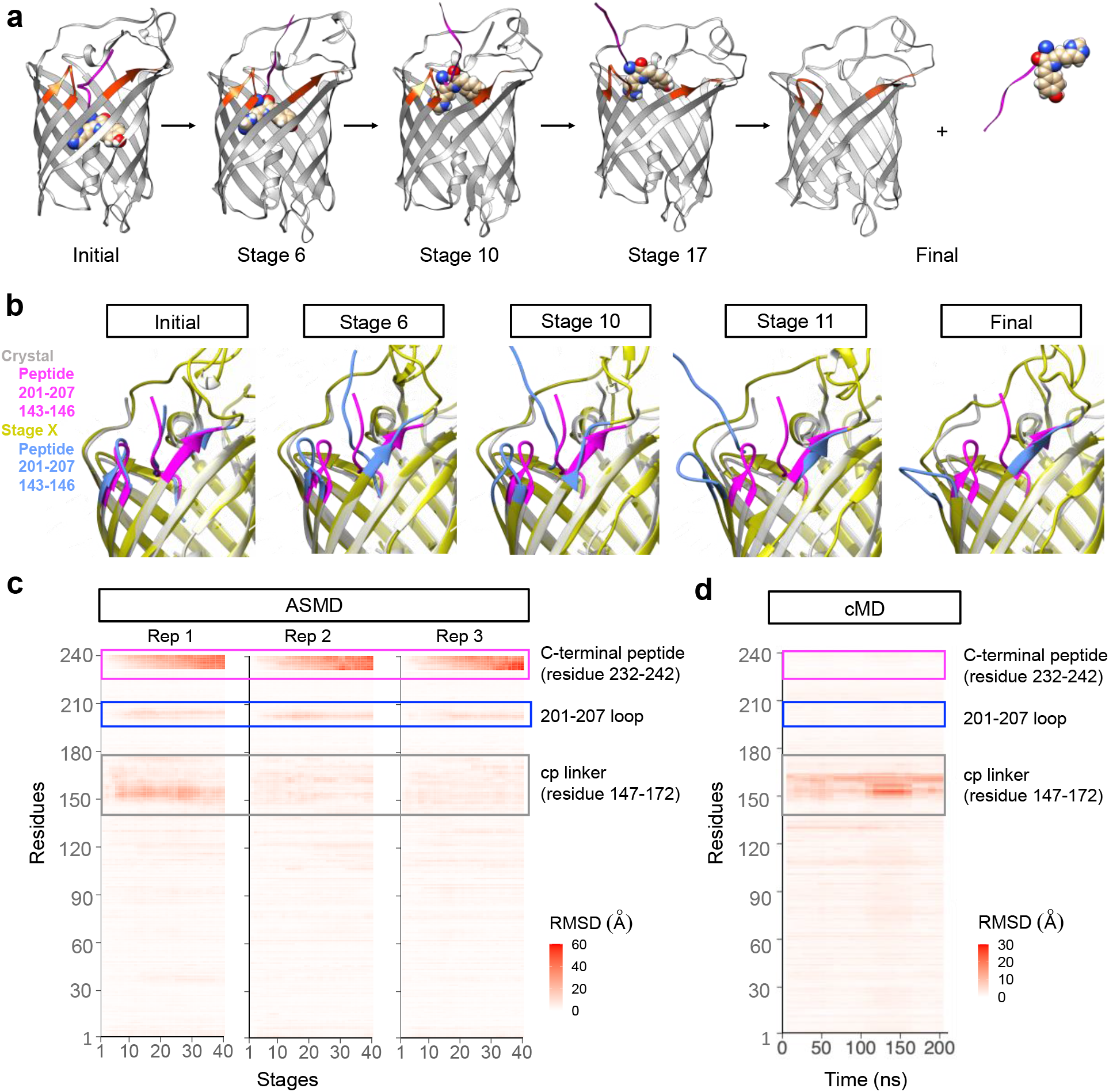
Molecular dynamic simulation on PhoCl1 dissociation process. (**a**) Representation of the simulated dissociation process. Photo-converted PhoCl1 is shown in grey, dissociated peptide fragment is shown as magenta ribbon for the peptide portion and spheres for the chromophore. Residues within and near the 201-207 loop are highlighted in orange. (**b**) Structure alignment of the crystal structure of the PhoCl1 red state and the simulated structure from different stages of ASMD replication 3 (Rep 3). Crystal structure of the PhoCl1 red state is shown in silver, with residues within and near the 201-207 loop highlighted in magenta. Simulated structures from different stages are shown in yellow, with residues within and near the 201-207 loop highlighted in blue. (**c**) RMSD heatmaps of 3 ASMD replications over stages with the initial stage as reference. The RMSD values were colored according to the scale bar. The C-terminal peptide is enclosed in a magenta box, the 201-207 loop is enclosed in a blue box and the cp linker is enclosed in a grey box. Rep, replication. (**d**) RMSD heatmaps of cMD over stages with the initial stage as reference. The RMSD values were colored according to the scale bar.

Analysis of the RMSD values for all residues over time, with the initial stage as the reference, revealed that the C-terminal peptide showed the highest RMSD due to its physical displacement (**Fig. 2c**). The region with the second highest RMSD was the 201-207 loop, consistent with conformational changes observed in the experimental crystal structures. Inspection of the dynamic conformational changes during stages of dissociation reveal that the 201-207 loop “flips” out of the barrel, presumably to minimize steric interactions with the C-terminal peptide as it dissociates (**Fig. 2b**). In contrast, such flipping behaviour of the 201-207 loop was not observed during 210 ns of unrestrained cMD of the red state (**Fig. 2d**). These results demonstrate that the 201-207 loop undergoes a dissociation-dependent conformational change, rather than being intrinsically flexible, providing support for the conclusion that the 201-207 loop would be the key region of interest in our subsequent engineering.

### Engineering and characterization of PhoCl2 variants

In an effort to further improve the utility of PhoCl, we used a combination of structure-guided rational engineering and directed evolution starting from the template of PhoCl1. We first established a NanoLuc luciferase^24^-based assay^25^ for the screening of PhoCl variants libraries on the basis of dissociation rate and efficiency (i.e., extent of dissociation) (**Fig. 3a**). PhoCl with long flexible linkers at both the N- and C-termini was incorporated into the NanoLuc Binary Technology (NanoBiT) system^25^. We found that, upon photoconversion of PhoCl with 30 s of illumination by a 405 nm LED array, dissociation could be monitored by the decrease in bioluminescence intensity associated with PhoCl-mediated dissociation of NanoBiT into its Large BiT (LgBiT; 18kDa) and Small BiT (SmBiT; 11 amino acid peptide) fragments. To increase the solubility of the SmBiT fragment (fused to the small peptide fragment of PhoCl), we also fused the maltose binding protein (MBP) to the C-terminus of the construct. Using this assay, we observed a −39% change in bioluminescence at 460 nm for PhoCl1 upon illumination (**Supplementary Fig. 3**).

**Figure 3.**
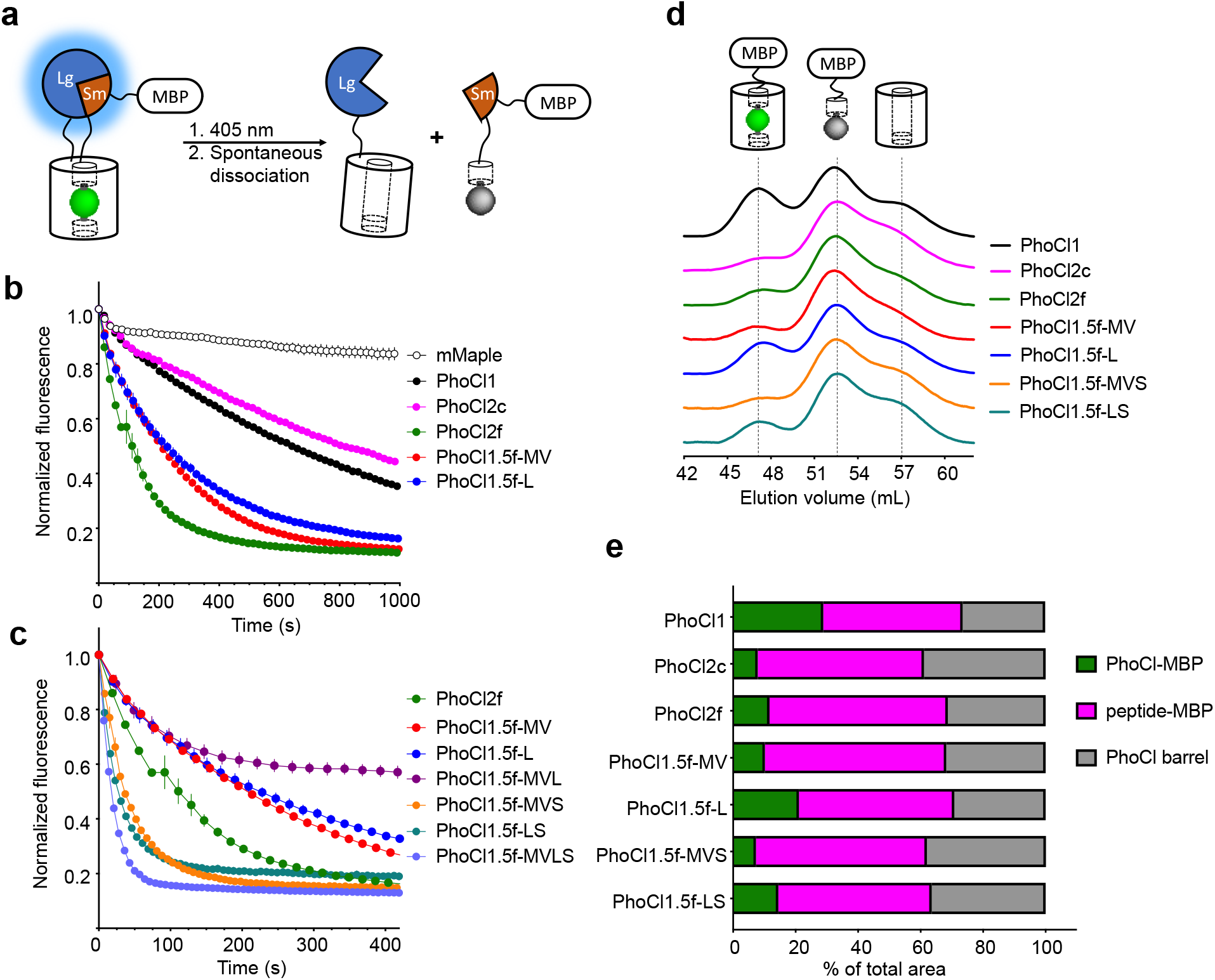
Library screening assay and characterization of PhoCl2 variants. (**a**) Schematic of the PhoCl screening strategy design. (**b)** Dissociation kinetics, as determined by loss of red fluorescence after photoconversion. The red fluorescence was monitored immediately following 15 s illumination with 405 nm light (LED array, 3.46 mW/mm2). Values are means ± SEM (*n* = 3 independent experiments). One phase decay fit was used to determine the dissociation half time, *t*_1/2_. *R*-squared values for fits range from 0.9830 to 0.9975. (**c**) Dissociation kinetics of PhoCl2f and PhoCl1.5f variants with the combinations of key mutations. (**d**) Gel-filtration chromatography (GFC) analysis of maltose binding protein (MBP) fused PhoCl variants. All proteins were tested at the same concentration (2 mg/mL) and illumination conditions (15 s LED array illumination, 3.46 mW/mm^2^). Partial photoconverted sample was loaded on the HiPrep 16/60 Sephacry S-100 column immediately after illumination. The intact PhoCl-MBP fusion (75 kDa) was eluted first from the column at a volume of ∼ 47 mL. This was followed by the cleaved peptide-MBP fusion (45 kDa) at ∼ 52 mL and then the cleaved empty barrel (30 kDa) at a volume of ∼ 57 mL. The identity of the fragments associated with each peak in the eluent were confirmed through SDS-PAGE analysis (in **Supplementary Fig. 8c**). (**e**) Summary of peak area ratios from the GFC analysis. Baseline correction, multi-peak fitting, and peak area integration were performed using the OriginLab software. A Gaussian fit model was used in peak fitting with three fixed widths curves corresponding to the three elution peaks. *R*-squared values for peak fits range from 0.9974 to 0.9993.

Photoconvertible protein mMaple^30^ (the progenitor of PhoCl) inserted into NanoBiT was used as a negative control for this assay. When using this negative control, we also observed a substantial decrease in apparent bioluminescence (−26% at 460 nm). We rationalize this decreased bioluminescence as a result of the change in Bioluminescence Resonance Energy Transfer^31^ (BRET) efficiency as the chromophore is converted from green to red. More specifically, BRET to the green state results in sensitized emission that overlaps with the emission of NanoLuc itself. BRET to the red state results in sensitized emission at a longer wavelength peak that does not overlap with that of NanoLuc. This explanation is consistent with the emerging bioluminescent peak at 583 nm corresponding to the mMaple red state (**Supplementary Fig. 3b)**.

Starting from the template encoding PhoCl1, two error-prone PCR libraries and 32 site-directed mutagenesis libraries were created. The residues involved in either hydrogen bonding or hydrophobic contacts with the chromophore and dissociable peptide, including the residues of the 201-207 loop (Ile3, Asp5, Phe7, Lys8, Phe11, Gly14, Tyr15, Ile35, Met37, Glu38, Gly39, Asp40, Phe42, Arg77, Asp78, Gly79, Val80, Met87, Tyr117, His118, Val120, His122, Leu138, Tyr139, Val143, Ala144, Arg145, Asn146, Met162, Asp177, Met178, Gly201, Lys202, Pro203, Phe204, Glu205, Gly206, Ile207, Gln208, Ile210, Thr229, Ala230, Thr239 and Lys240) (**Supplementary Fig. 4** and **Supplementary Table 2**), were randomised by site-directed mutagenesis to introduce a codon (NNK) encoding all 20 amino acids into the corresponding position of the gene. We did not include the residues that are directly hydrogen bonded to the chromophore, in order to avoid disruption of chromophore formation. Green fluorescent clones were picked from libraries expressed in colonies of *Escherichia coli* on Petri dishes, and bacteria were cultured overnight in small volumes of liquid culture, and the bioluminescence assay was applied in the context of the cell culture lysate. The top ∼2% variants exhibiting large bioluminescence intensity decreases within 5 min after photoconversion were picked for DNA sequencing and also served as templates for subsequent engineering. We observed that most colonies from the site-directed mutagenesis libraries exhibited decreased green fluorescence, likely due to decreased efficiency of chromophore formation. We reasoned that poor chromophore formation was undesirable because proteins without proper chromophore cannot be photoconverted and undergo dissociation, and so we biased our picking towards brighter green fluorescent clones. Notably, this approach may have missed variants that exhibited reduced green fluorescence due to a reduced quantum yield of fluorescence and that may have retained efficient chromophore formation and photocleavage capability.

**Figure 4.**
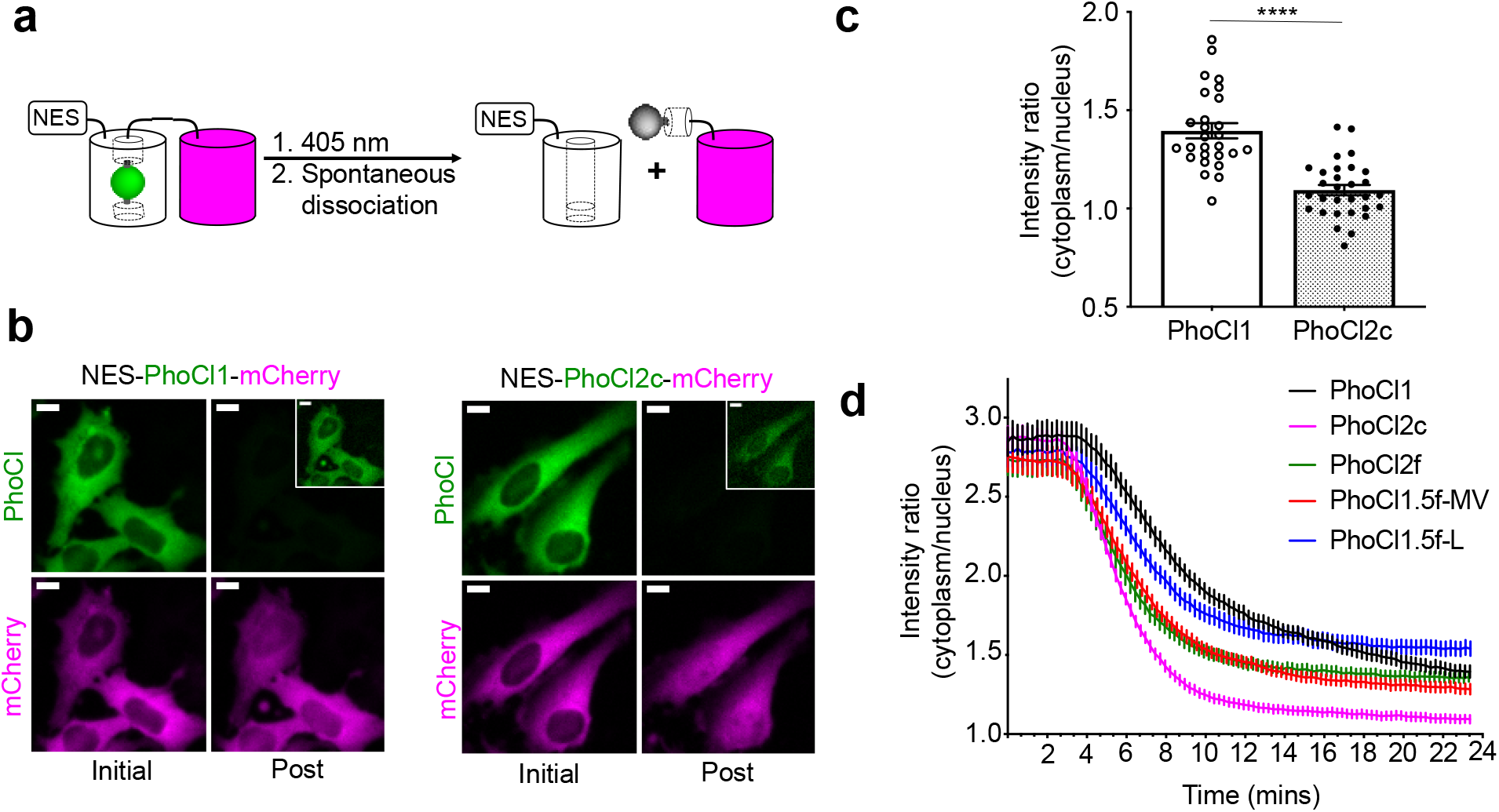
Optogenetic manipulation of protein translocation in HeLa cells. (**a**) Schematic of the NES-PhoCl-mCherry photocleavage. (**b**) Representative images of HeLa cells expressing NES-PhoCl-mCherry before and after (15 min) photoconversion. Conversion was performed with 10 s violet light pulses (395/40 nm, 2 mW/mm^2^) every 15 s for 6 mins. Inset is the same image with 10× increased contrast. Scale bar, 10 µm. (**c**) Red fluorescence intensity ratios of cytoplasm to nucleus at 15 min after photoconversion. Ratios were calculated for single cells. Values are means ± SEM (*n* = 27 cells of PhoCl1, and *n* = 30 cells of PhoCl2c). \*\*\*\**P* < 0.0001 by unpaired two-tailed *t* test (*t* (55) = 6.606). (**d**) Red fluorescence intensity localization ratios of cytoplasm to nucleus versus time. Values are means ± SEM (*n* = 27 cells of PhoCl1 and PhoCl1.5f-L, and *n* = 30 cells of the other variants).

Following extensive screening and directed evolution using the bioluminescence assay, we identified a number of specific mutations (Val143Met, Ala144Val, Lys202Leu and Ile207Ser) in or near the 201-207 loop that seemed to be associated with faster dissociation kinetics (**Fig. 3b**,**c, Supplementary Figs. 5, 6, 7**, and **Supplementary Table 3**). To investigate the dissociation kinetics in greater detail, we turned to our previously reported assay that relies on the loss of red fluorescence after photoconversion^5^. The red fluorescence intensity is monitored immediately following 15 s illumination with 405 nm light (LED array, 3.46 mW/mm^2^) (**Fig. 3b**,**c** and **Supplementary Fig. 8a**,**b**). Consistent with our bioluminescence screening assay, we found that variants with mutations in or near the 201-207 loop (PhoCl1.5f-MV, PhoCl1.5f-L, and PhoCl2f), exhibited faster dissociation rates (*t*_1/2_ = 76 - 140 s) than PhoCl1 (*t*_1/2_ = 570 s) (**Supplementary Table 4**). We next combined these key mutations to explore whether they might have synergetic beneficial effects on the rate of dissociation (**Fig. 3c** and **Supplementary Fig. 8b**). Testing a variety of combinations led to the PhoCl1.5f-MVS variant (with Val143Met, Ala144Val and Ile207Ser), the PhoCl1.5f-LS variant (with Lys202Leu and Ile207Ser), and the PhoCl1.5f-MVLS variant (with all four key mutations), all of which exhibited substantially faster dissociation rates (*t*_1/2_ = 15 - 30 s). The PhoCl1.5f-MVL variant (**Fig. 3c**), with a combination of Val143Met, Ala144Val and Lys202Leu (without Ile207Ser), has unexpectedly poor dissociation efficiency (with ∼60% red fluorescence remaining as shown in **Fig. 3c**), illustrating the delicate nature the dissociation process and the importance of empirical screening. All the variants with combined mutations were found to have relatively dim fluorescence brightness (in **Supplementary Table 4**) possibly due to a decreased efficiency of chromophore formation. The PhoCl1.5f-MVLS variant also has particularly poor expression, possibly indicative a poor folding efficiency, and thus we were unable to purify sufficient protein for a full characterization. We ultimately determined that the variant with mutations Lys49Gln, Tyr117Cys, Arg145Lys, Asn146Thr and Ile207Ser (designated as PhoCl2f; *t*_1/2_ = 76 s) represented the best compromise of dissociation rate, fluorescence brightness, and protein expression level.

**Figure 5.**
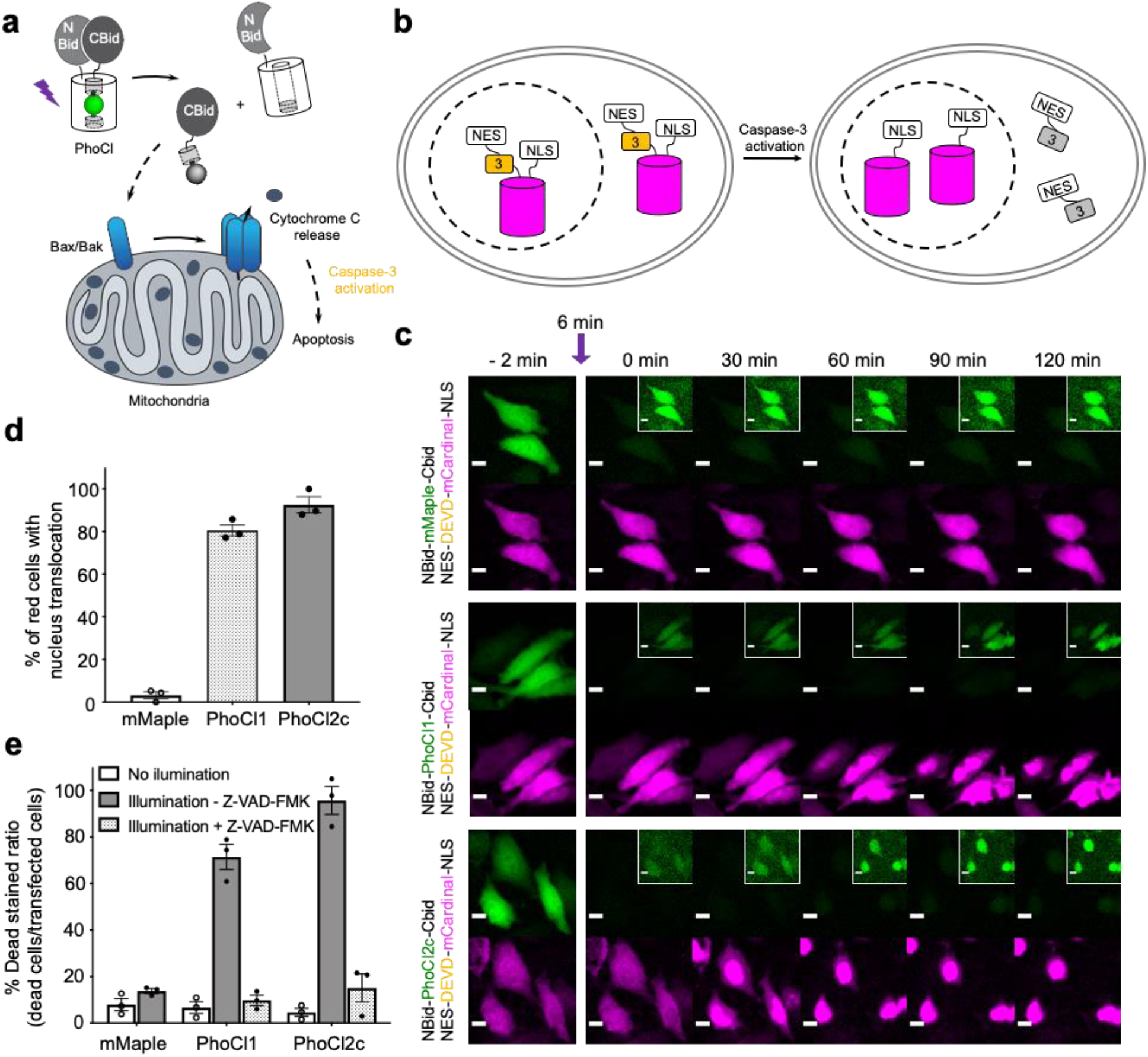
Optogenetic manipulation of cell apoptosis via PhoCl. (**a**) Schematics of optogenetic activation of apoptosis with a PhoCl-Bid construct. (**b**) Schematics of apoptosis reporter: NES-DEVD-mCardinal-NLS used in this experiment. (**c**) Transient transfected HeLa cells co-expressing NBid-mMaple-CBid or NBid-PhoCl-CBid with caspase-3 reporter. Cells were illuminated with 10 s violet light pulses (395/40 nm, 2 mW/mm^2^) every 15 s for 6 mins. Inset is the same image with 10× increased contrast. Scale bar, 10 µm. (**d**) Summary data of caspase-3 reporter translocation for cells expressing constructs described in **c**. Values are means ± SEM (*n* = 3 cell cultures of each variant. A total of 87 cells for mMaple, 89 cells for PhoCl1 and 113 cells for PhoCl2c). (**e**) Summary data of cell viability assay. Photoconversion was performed with 30 s pulse light (LED array, 3.46 mW/mm^2^). Ratios are calculated by determining the ratio of DEAD stained cells to green transfected cells. Values are means ± SEM (*n* = 3 cell cultures of each variant). A total of 321 cells for mMaple no illumination, 301 cells for mMaple illumination - Z-VAD-FMK, 316 cells for PhoCl1 no illumination, 328 cells for PhoCl1 illumination - Z-VAD-FMK, 326 cells for PhoCl1 illumination + Z-VAD-FMK, 325 cells for PhoCl2c no illumination, 337 cells for PhoCl2c illumination - Z-VAD-FMK and 318 cells for PhoCl2c illumination + Z-VAD-FMK.

In parallel to the engineering of faster-dissociating variants, we also identified PhoCl2c (with mutations Lys49Gln, Cys99Gly, Tyr117Cys, Arg145Lys and Asn146Thr) with enhanced dissociation efficiency. To characterize the dissociation efficiency of new PhoCl variants, we used a modified version of our NanoLuc-based bioluminescence screening assay in which we compared the bioluminescence emission spectrum before illumination and 30 minutes after illumination (**Supplementary Fig. 3** and **Supplementary Table 5**). Most PhoCl2 variants exhibited substantial decreases in bioluminescence intensity after illumination, consistent with our expectations (**Supplementary Fig. 3a**). PhoCl2c and PhoCl1.5f-MVS variants showed the largest decrease around 60%, which indicates that these 2 variants have the best dissociation efficiency. To investigate the dissociation efficiency in greater detail with an orthogonal method, we turned to gel filtration chromatography (GFC) assay combined with analysis of the eluted fractions by SDS-PAGE analysis (**Fig. 3d**,**e** and **Supplementary Fig. 8c**). This analysis demonstrated that the PhoCl2c-MBP fusion had higher dissociation efficiency with less remaining intact protein (7.9% intact protein) than the PhoCl1-MBP fusion (28.9% intact protein). Even with the molecular structure of PhoCl1 in hand, it remains unclear to us why PhoCl2c exhibits higher dissociation efficiency. For example, one of the key mutations of PhoCl2c, Cys99Gly, is located on the opposite end of the barrel from the dissociable peptide. PhoCl1.5f-MVS variant has a similarly high dissociation efficiency (7.3% intact protein) as PhoCl2c, consistent with the bioluminescence analysis, but again the relatively poor chromophore formation of PhoCl1.5f-MVS will likely limit its further application. Taken together, we successfully engineered two second-generation PhoCl variants: PhoCl2c with higher dissociation contrast ratio, and PhoCl2f with faster dissociation rate.

### Optogenetic control of protein localization by PhoCl2 variants in HeLa cells

The dissociation rate and efficiency of the PhoCl2 variants was further characterized in transiently transfected HeLa cells using our previously reported assay for manipulating subcellular protein localization through PhoCl-dependent cleavage of a nuclear exclusion sequence (NES)^5,32^ (**Fig. 4a**). Photocleavage leads to redistribution of a red FP (mCherry) between the cytoplasm and nucleus. Relative to PhoCl1-mCherry, PhoCl2c-mCherry fusion exhibited a more even red fluorescence redistribution (between the cytoplasm and nucleus) following photocleavage to remove the NES (**Fig. 4b**). Following protein redistribution after photocleavage, the cytoplasm-to-nucleus intensity ratio for PhoCl2c-mCherry is lower than the ratio for PhoCl1-mCherry (**Fig. 4c**).

The kinetics of photoconversion was determined by loss of green fluorescence, and dissociation was characterized by red fluorescence redistribution in HeLa cells expressing NES-PhoCl-mCherry (**Fig. 4d** and **Supplementary Table 6**). PhoCl1.5f-MV, PhoCl1.5f-L and PhoCl2f exhibited faster mCherry redistribution and similar dissociation efficiency (in **Supplementary Fig. 9**) compared to PhoCl1, which is consistent with our *in vitro* protein characterization results. However, we did find that three other 1.5f variants (PhoCl1.5f-MVS, PhoCl1.5f-LS and PhoCl1.5f-MVLS) did not dissociate as efficiently in cells (refer to **Supplementary Fig. 9**) as expected, possibly due to poor efficiency of chromophore formation. These results collectively establish the improved characteristics of PhoCl2c and PhoCl2f in cellular assays for optogenetic control.

### Optogenetic manipulation of cell apoptosis using PhoCl2c

We next attempted to explore possible optogenetic manipulation of cell apoptosis through the use of photocleavage to activate BH3 interacting-domain death agonist (Bid). Bid is a pro-apoptotic protein belonged to Bcl-2 family. Following cleavage by caspase-8 during apoptosis, Bid translocates to the mitochondria, where it then induces an increase in outer mitochondria membrane permeability and cytochrome c release^33,34^. To achieve the optogenetic control of apoptosis, we engineered a photoactivatable Bid by inserting PhoCl into the middle of Bid at the position of the caspase-8 cut site (LQTDG)^35^. We reasoned that photoinduced dissociation of C-terminal domain (CBid) from its autoinhibitory N-terminal domain (NBid) would induce the downstream apoptosis pathway (**Fig. 5a**). We monitored the downstream caspase-3 activity using a NES-DEVD-mCardinal-NLS reporter. The cleavage of DEVD linker of the fusion protein by caspase-3 activation results in the translocation of red FP mCardinal into the nucleus (**Fig. 5b**). Caspase-3 activation tended to be observed at around 30 min after conversion for the NBid-PhoCl2c-CBid and around 50 min after conversion for NBid-PhoCl1-CBid (**Fig. 5c, Supplementary Fig. 10a**,**b** and **Supplementary Movies 5-7**). Cell shrinkage and rounding followed by FP translocation were observed at the end of 2 hours of imaging photoconverted cells. We used the non-dissociable mMaple insertion construct as negative control as it is not expected to dissociate and induce FP translocation following illumination (**Fig. 5d**). Cells expressing NBid-mMaple-CBid or NBid-PhoCl-CBid variants were illuminated and then cells were analyzed using the DEAD cell viability assay (shown in **Fig. 5e** and **Supplementary Fig. 10c-e**). The dye in DEAD stain is ethidium homodimer-1 which is used to indicate loss of cell plasma membrane integrity. For cells expressing NBid-mMaple-CBid, no significant illumination-dependent difference in cell viability was observed. In contrast, illumination of cells expressing NBid-PhoCl1-CBid exhibited significantly higher cell death ratios than non-illuminated cells. A further enhanced death rate was observed with PhoCl2c due to its higher dissociation efficiency. Treatment with the cell permeable pan-caspase inhibitor Z-VAD-FMK^36^ effectively reversed this effect, illustrating that cell death was indeed due to caspase activation during apoptosis (shown in **Supplementary Fig. 10d**,**e**). In summary, we have designed a novel PhoCl-based optogenetic tool for induction of cell apoptosis pathway. Compared to NBid-PhoCl1-CBid, NBid-PhoCl2c-CBid exhibited more efficient light-dependent induction of apoptosis, providing support for the conclusion that the improved properties (as characterized with purified protein) have been retained for the protein expressed in mammalian cells.

## Discussion

In this work we have described the development of two improved versions of the PhoCl photocleavable protein: the PhoCl2c variant with enhanced dissociation efficiency, and the PhoCl2f variant with faster dissociation rate. This advance was made possible by the determination of the X-ray crystal structure of the empty barrel fragment which revealed substantial conformational changes in the 201-207 loop. During library screening, we discovered four key mutations found within or near 201-207 loop which had a substantial and favorable effect on the PhoCl dissociation rate: Lys202Leu and Ile207Ser in the loop and the combination of Val143Met and Ala144Val on the adjacent β-strand 11 (**Supplementary Fig. 7**). Overall, these mutations are clustered in close vicinity to where the C-terminal peptide emerges from the intact protein. It is reasonable to postulated that mutations in this region would affect the efficiency and rate of peptide dissociation. At this point the mechanisms by which these mutations are exerting their observed influence on peptide dissociation is unclear. Two possible mechanisms are: 1) Destabilization of the red state, leading to a decreased energy barrier for peptide dissociation and faster kinetics; and 2) Increased stabilization of the empty barrel which would likely have little effect on the kinetics, but may lead to a greater overall efficiency of dissociation.

The critical importance of mutations in the loop region was made apparent by the bioluminescence assay which monitored the rate of dissociation. However, this same assay did reveal some unexpected differences in BRET efficiency between different PhoCl variants (refer to **Supplementary Fig. 3b** and **Supplementary Table 5**). As with FRET^37^, BRET efficiency strongly depends on the interchromophore distance and orientation. With the same luciferase donor and linkers, the BRET ratio is also related to extinction coefficient and quantum yield of PhoCl acceptor. After photoconversion, the BRET signal should decrease or even disappear as a result of the photocleavage of the green chromophore. Compared to PhoCl1, most of the variants showed decreased BRET efficiencies (that is, higher ratios of NanoLuc emission at 460 nm to PhoCl emission at 505 nm), which can be explained by the diminished extinction coefficients and reduced quantum yields. For these variants with low BRET efficiency, bioluminescence intensity at 460 nm decreased upon illumination and photocleavage, as expected. However, for reasons that remain unclear to us, PhoCl1.5f-MV and PhoCl1.5f-L exhibited substantially higher BRET efficiency than other variants. Moreover, the luminescence of these 2 variants increased after photoconversion which is contrary to what we observed with the other variants. We speculate that Lys202Leu in PhoCl1.5f-L, and the combination of Val143Met and Ala144Val in PhoCl1.5f-MV, may have introduced stabilizing interactions between NanoLuc and PhoCl that resulted in a more favorable distance or orientation for increased BRET efficiency. The high BRET ratio still existed in partial converted protein fusion after illumination. Hence, the luminescence signal at 460 nm was observed to increase as a result of the reduced BRET efficiency.

An important factor that is likely contributing to the improved performance of all of the new variants reported in this work are the increased extinction coefficients at 405 nm (**Supplementary Table 4** and **Supplementary Fig. 11**), which would be expected to lead to more efficient absorption of 405 nm light and more efficient photoconversion. This higher absorbance at 405 nm may help to explain why the PhoCl2f and PhoCl1.5f variants, which were selected based due to their faster kinetics of dissociation, also exhibited higher dissociation efficiency than PhoCl1 by GFC analysis (**Fig. 3d**,**e** and **Supplementary Fig. 8c**). In our mammalian cell-based assays, the PhoCl2c constructs exhibited faster redistribution (**Fig. 4d** and **Supplementary Table 6**), and faster induction of cell apoptosis, than the corresponding PhoCl1 constructs (**Fig. 5c** and **Supplementary Fig. 10a**,**b**). These results are at odds with our *in vitro* kinetics analysis which demonstrated that PhoCl2c was slightly slower at dissociating than PhoCl1 (**Fig. 3b**). However, suggest that this discrepancy can be explained by the PhoCl2c’s higher photoconversion efficiency due to higher absorbance of 405 nm light. This result indicates that both the efficiency of photoconversion, and the kinetics of dissociation, are contributing to the effective rate of release in cells.

To summarize, we described the structure-guided engineering of the second generation of PhoCl variants (PhoCl2) with improved dissociation extent (PhoCl2c) and rate (PhoCl2f) by using the luciferase-based screening assay. We expect that this high-throughput screening method could potentially be used to engineer other improved photosensory domains in optogenetic actuators that undergo light-induced heterodimer to monomer transitions (e.g., LOVTRAP^38^ and PhyB-PIF^16^). Finally, we anticipate that PhoCl2 variants will prove useful to in an increased range of optogenetic applications due to their faster and more efficient photo-induced cleavage and dissociation.

## Methods

### General methods and materials

All synthetic DNA oligonucleotides were purchased from Integrated DNA Technologies (IDT). Primer sequences are provided in **Supplementary Table 7**. Plasmid constructions were performed with standard restriction enzyme cloning or In-fusion HD Cloning kit (Takara Bio USA) according to the manufacturer’s protocol. Q5 high-fidelity DNA polymerase (New England BioLabs) was used for standard polymerase chain reaction (PCR) amplification, and Taq DNA polymerase (New England BioLabs) was used for error-prone PCR. Restriction enzymes and T4 ligase were purchased from Thermo Fisher Scientific. Site-directed mutagenesis was performed using QuikChange Lightening kit (Agilent). Transformation of bacterial cells with plasmid DNA was performed by electroporation using a MicroPulser Electroporator (Bio-Rad). GeneJET gel extraction kit and plasmid miniprep kit (Thermo Fisher Scientific) were used for DNA extraction. DNA sequencing was performed by the University of Alberta Molecular Biology Service Unit.

### Library constructions, mutagenesis and screening

For constructing the PhoCl screening vector, the *NanoBiT* template was ordered as a synthetic gBlock from IDT. The *NanoBiT* gene sequence was generated using Nluc gene (GenBank: JQ437370.1) as the template, and combining the amino acid substitutions reported in literature^25^. A gene encoding LgBiT-PhoCl-SmBiT containing 5’ *KpnI* and 3’ *XbaI* restriction sites at the ends of *PhoCl* was made by overlap extension PCR. Each terminus of *PhoCl* and split *NanoBiT* fragments were joined by linker sequences reported in a previous study^25^. The gene fragments encoding LgBiT-PhoCl-SmBiT (with 5’ *XhoI* and 3’ *EcoRI* restriction sites) and maltose binding protein (MBP) (with 5’ *EcoRI* and 3’ *HindIII* restriction sites) were digested and inserted between the *XhoI* and *HindIII* sites of pBAD/HisB vector by three-way ligation thus yielding the PhoCl screening construct of *XhoI*-*LgBiT*-*KpnI*-*PhoCl*-*XbaI*-*SmBiT*-*EcoRI*-*MBP*-*HindIII*.

The *PhoCl* libraries were generated by error-prone PCR or site-directed mutagenesis with degenerate codons at the targeted sites. *E. coli* strain DH10B (Thermo Fisher Scientific) electrocompetent cells were transformed with the gene libraries and grown on lysogeny broth (LB) agar plates supplemented with 100 µg/mL ampicillin and 0.02% L-arabinose. For libraries generated by error-prone PCR, approximately 10,000 colonies were screened in a given round. The colonies exhibiting the brightest green fluorescence (top ∼ 2%), as determined using a custom-built colony screener^5^ (for green channel: 470/40 nm excitation and 510/20 nm emission), were picked and cultured in 1 mL medium in 96-well deep block plates (Thermo Fisher Scientific). For libraries generated by randomization of one codon, 190 colonies (approximately six-fold the theoretical number of gene variants generated from an NNK codon, where N = A, G, C, T and K = G, T) were picked randomly and screened. For libraries generated by randomization of a combination of two codons, 570 green fluorescent colonies (approximately 50% of the theoretical number of gene variants) were picked and screened.

For each cultured variant, protein was extracted using 50 µL 0.5% of *n*-octyl-β-D-thioglucopyranoside in Tris buffered saline (TBS, pH 8) and then further diluted 1:4 in PBS buffer (pH 7.4). Photoconversion was performed in transparent microplates (Nunc-Thermo Fisher Scientific; Mfr.No. 265302) under a 405 nm LED flood array (Loctite) with an intensity of 3.46 mW/mm^5^. Diluted samples (100 µL), with and without photoconversion, were tested on 96-well plates (Thermo Fisher Scientific) at 5 min after photoconversion. To each well was added 10 µL luciferase substrate furimazine in PBS solution (0.1 mg/mL), and the luminescence was measured immediately using a Cytation 5 plate reader (BioTek). The top 2% variants with bright bioluminescence and large intensity decrease (> 60% decrease) were picked for sequencing and used as templates for the next evolution. A total of 34 libraries with 46 targeted amino acids were screened for the identification of the PhoCl2c and PhoCl2f variants.

### Protein purification and *in vitro* characterization

To purify protein for crystallization, the gene expressing PhoCl with a 6× His tag at the C-terminal in pET-28a vector was used to transform *E. coli* strain BL21 (DE3) pLysS (Promega). A single colony was picked to inoculate a 5 mL starting culture that was allowed to grow overnight at 37 °C with 225 rpm before being diluted into large cultures (1 liter for PhoCl green state and red state; 10 liter for PhoCl empty barrel) of LB media supplemented with 100 µg/mL ampicillin. The cultures were allowed to grow at 37 °C to an OD_600_ of 0.5, and then induced with IPTG (0.1 mM) and grown overnight at 28 °C. The *E. coli* cells were harvested by centrifuge and lysed by sonication. Protein was purified by Ni-NTA chromatography (G-Biosciences). For PhoCl empty barrel purification, the intact 6× His tag fused PhoCl protein was captured by Ni-NTA agarose and was photoconverted on the column with 10 × 15 s illumination (LED array, 3.46 mW/mm^2^), and the eluent containing the PhoCl empty barrel was collected.

To prepare the proteins for *in vitro* characterizations of PhoCl variants, the *E. coli* strain DH10B were transformed with the pBAD/HisB vector encoding the gene of interest. Cultures were induced with L-arabinose (0.2%) and allowed to grow 24 h at 30 °C. Protein were purified by Ni-NTA chromatography. Molar extinction coefficients (ε) at 488 nm were determined by the alkali denaturation method^39^ and then used as reference ε for the 405 nm. Quantum yields for PhoCl variants were measured using purified EGFP as the reference standard^40^. Photoconversion was performed with the 405 nm LED flood array. The absorbance spectra were acquired with a DU-800 UV-vis spectrophotometer (Beckman). The fluorescence spectra were acquired with a Safire2 plate reader (Tecan) and bioluminescence spectrums were acquired with a Cytation 5 plate reader (BioTek). The gel filtration was performed with a HiPrep 16/60 Sephacryl S-100 column on an AKTA chromatography system (GE Healthcare), and the size of the proteins in each eluted fraction were verified by SDS-PAGE.

### Protein crystallization and X-ray data collection

The green state of PhoCl1 and PhoCl1 empty barrel recombinant proteins were further purified with size exclusion chromatography using SD200 column (GE Healthcare) and buffer exchanged to 20 mM Tris pH 7.4, 150 mM NaCl. Initial crystallization trials were carried out with 384-well plate via sitting drop vapor diffusion against commercially available sparse matrix screens (Hampton Research and Molecular Dimensions) at room temperature. The green state of PhoCl1 was crystallized in 0.1 M MIB buffer pH 6.0, 25% w/v PEG 1500. Crystals of the red state of PhoCl1 was generated from the PhoCl1 green crystals through photoconversion with the violet light (15 s illumination with 405 nm LED array, 3.46 mW/mm^2^). The PhoCl1 empty barrel crystals were observed in 0.056 M NaH_2_PO_4_·H2O, 1.344 M K_2_HPO_4_, pH 8.2. All the crystals were cryoprotected with the reservoir condition in supplement with 15-20% glycerol and flash frozen in liquid nitrogen. X-ray diffraction dataset were collected at Canadian Light Source CMCF-ID beamline. All the datasets were processed and scaled with XDS package^41^; the data collection details and statistics are summarized in **Supplementary Table 1**.

### Structure determination and refinement

The structure of the green state of PhoCl1 was solved by molecular replacement using mTFP1 (PDB ID: 2HQK) as search model^42,43^. The red state of PhoCl1 and PhoCl1 empty barrel structure determination were carried out using the solved the green state of PhoCl1 as template. The manual model building and refinement were performed with Coot^44^ and PHENIX^45^ refine. The CR8 and IEY were used as the chromophore ligands for the green state and the red state of PhoCl1 respectively. In the PhoCl1 red crystal, the chromophore turned out to be partially converted, therefore two of the six monomers in the asymmetric unit with high occupancy for IEY were modeled in red form, and the other four monomers were built in green form.

The PhoCl1 green form was solved to 2.10 Å with a *R*_*Work*_ factor of 0.1824 and *R*_*Free*_ factor of 0.2162 in P2_1_2_1_2_1_ spacegroup. The Photoconversion of the PhoCl1 green crystal induced the packing of the crystal and the PhoCl1 red model was refined to 2.30 Å with a *R*_*Work*_ factor of 0.2186 and *R*_*Free*_ factor of 0.2638 in P1 spacegroup. The PhoCl1 empty barrel was determined to 2.82 Å in P2_1_ spacegroup with anisotropical diffraction; the dataset was processed with the Diffraction Anisotropy Server^46^ and refined with a *R*_*Work*_ factor of 0.2587 and *R*_*Free*_ factor of 0.3029.

All the structures showed favorable stereochemistry and exhibited good distribution of dihedral angles on a Ramachandran plot. Detailed refinement statistics are summarized in **Supplementary Table 1**. Structure figures were generated using the molecular visualization PyMOL package (version 1.8)^47^ and UCSF Chimera (production version 1.13.1)^48^. Schematic diagram of the dissociable peptide in the PhoCl1 red state was generated using LIGPLOT (version 4.5.3)^49^.

### Molecular dynamic (MD) simulation

To prepare the starting structure for MD, the red state chromophore in the protonated state, designated as “RCP”, was generated in Avogadro (version 1.2.0)^50^, and parameterized using R.E.D. Server Development 2.0 (Ref. 51). The circular cp linker (residues 147-172), which was not resolved in the crystal structure of the photo-converted PhoCl1, was modelled in Rosetta (version 2018.33.60351) using the kinematic closure (KIC) with fragment protocol^52^. Briefly, 10,000 models were generated and the structure with the largest negative score was selected to represent the starting conformation of the cp linker. The experimental structure of the PhoCl1 red state structure, edited to include the parameterized RCP chromophore and the cp linker, was subsequently solvated in LEaP (AmberTools19) with explicit water (TIP3P) and 1 sodium counterion in an octagon periodic boundary condition with each side no closer than 30 Å away from the protein.

All MD simulations are performed with Amber18 with GPU parallelization on Cedar cluster, Compute Canada. The system was minimized, heated to 303.15 K and equilibrated with decreasing restraints for the protein. An unconstrained production simulation (cMD) was carried out at 303.15 K for 210 ns under isothermal–isobaric (NPT) ensemble. The restart file at the end of the cMD served as the initial coordinate of the adaptive steered MD (ASMD). The distance between the centres-of-mass (COMs) for the barrel (residues 1-231) and the C-terminal peptide (residues 232-242) were subjected to a harmonic constraint (rk2 = 18) increasing by 40 Å over 40 ns, without specifying the direction of dissociation. Such process was divided into 40 stages, where the distance between the COMs increased by 1 Å over 1 ns in each stage with 100 replications. The average of potential of mean force (PMF) was calculated by Jarzynski Equality^53^ after all replicates had completed. The replicate closest to the average was chosen to represent the trajectory of that stage whose restart file served as the initial coordinate of the next stage. To investigate whether the conformation of the cp linker affects the dissociation pathway, the same ASMD simulation was repeated twice restarting from 150 ns and 200 ns of the cMD, respectively. The trajectories of the chosen replicate from each stage were concatenated and aligned with the program CPPTRAJ (AmberTools19)^54^. Free energy along the dissociation process was calculated using molecular mechanics/Poisson-Boltzmann surface area (MM/PBSA) method quasi-harmonic entropy calculation^55^ with 20 snapshots from each stage. RMSD analysis and movie preparation were performed with VMD (version 1.9.4a31)^56^ software.

### PhoCl expression vectors and cell transfection

The previously reported pcDNA-NES-PhoCl-mCherry (Addgene #87690) plasmid was used as the backbone vector for the protein translocation experiments. A *HindIII* restriction site was introduced at the 3’ end of the NES sequence in the vector with the QuikChange Lightening kit. The DNA fragments encoding the PhoCl variants (with 5’ *HindIII* and 3’ *KpnI* restriction sites) were amplified via PCR and inserted into the appropriately digested NES-PhoCl-mCherry vector. For the construction of PhoCl inserted Bid, human *Bid* template was ordered as a synthetic gBlock from IDT. The DNA sequence was backtranslated from protein sequence BH3-interacting domain death agonist isoform 2 (GenBank: NP_001187.1) with codon optimization for *Homo sapiens* expression. The gene encoding NBid-PhoCl-CBid, 5’ *BamHI* and 3’ *KpnI* restriction sites at the ends of *PhoCl*, was made using overlap extension PCR to connect three gene fragments (i.e., NBid, PhoCl, and CBid). The assembled gene (with 5’ *NheI* and 3’ *XhoI* restriction sites) was digested and inserted between the *NheI* and *XhoI* sites of pcDNA 3.1(+), thus yielding the PhoCl inserted Bid expression vector.

To construct the caspase-3 reporter, DNA encoding the NES of mitogen-activated protein kinase kinase (MAPKK)^32^ and caspase-3 substrate (DEVD) was appended by PCR to the 5’ end of the gene encoding mCardinal, and DNA encoding the 3 × NLS of SV40 (Ref. 57) was appended to the 3’ end of fragment. The gene encoding NES-DEVD-mCardinal-NLS was then inserted between *NheI* and *XhoI* sites of pcDNA 3.1(+), thus yielding the caspase-3 reporter expression vector.

For expression of NES-PhoCl-mCherry, HeLa (ATCC CCL-2) cells at 70% confluency in 35-mm cell culture dishes (Corning) were transfected with 2 µg plasmid DNA and 4 µL Turbofect (Thermo Fisher Scientific) according to the manufacturer’s protocol. Imaging was performed at 24 h after transfection. For expression of PhoCl inserted Bid and caspase-3 reporter, HeLa cells were co-transfected with 1.75 µg NES-DEVD-mCardinal-NLS expression vectors and 0.25 µg PhoCl inserted Bid expression vectors mixed with 4 µL Turbofect. For expression of PhoCl inserted Bid for the DEAD viability assay, 1.75 µg pcDNA 3.1(+) empty vector and 0.25 µg PhoCl-Bid were used per transfection. The transfection was performed in 2 mL of serum-free DMEM for 2 h, after which the medium containing the transfection reagent and DNA mixture was replaced with 2 mL of complete medium supplemented with 10% fetal bovine serum (FBS; Sigma-Aldrich). Transfected cells were cultured 16 h before experiments.

### Cell photoconversion and imaging conditions

Before imaging, the medium was changed to HEPES-buffered Hanks Balanced Salt Solution (HHBSS; 20 mM HEPES). Wide-field fluorescence imaging was performed using a Nikon Eclipse Ti-E epifluorescence microscope equipped with a 75-W Nikon xenon lamp and a Photometrics QuantEM 512SC camera. The NIS-Elements AR package software was used for automatic instrument control and image acquisition. Cells were imaged with a 20× air objective lens (NA 0.75; Nikon) using the following filter sets: PhoCl green (490/15 nm excitation: ex; 510 nm dclp dichroic and 525/50 nm emission: em), mCherry and DEAD stain (543/10 nm ex, 565 nm dclp dichroic and 620/60 nm em), photoconversion (395/40 nm ex and 425 nm dclp dichroic), and mCardinal (605/50 nm ex, 635 nm dclp dichroic and 670/50 nm em).

For imaging of NES-PhoCl-mCherry protein translocation, the images were acquired in both the PhoCl green and mCherry channel every 15 s for 2 min before photoconversion. Photoconversion was performed with 10 s pulse illumination (2 mW/mm^2^) every 15 s for 6 min and each pulse of photoconversion light was followed by acquisition of PhoCl and mCherry fluorescence images. Imaging acquisition of green and red channels continued every 15 s for another 15 min after the photoconversion. Protein translocation was analyzed as the ratio of intensity of mCherry in the cytoplasm to the nucleus. Photoconversion efficiency was determined by the loss of green fluorescence intensity in the cytoplasm.

For imaging the caspase-3 activity triggered by photo-induced cell death with PhoCl inserted Bid, images were acquired in both the PhoCl green and mCardinal channel every 15 s for 2 min before photoconversion. Photoconversion was performed with 10 s pulse illumination (2 mW/mm^2^) every 15 s for 6 min and each pulse conversion light was followed by acquisition of PhoCl and mCardinal fluorescence images. Imaging acquisition of green and red channels continued every 30 s for 2 h after the photoconversion. Caspase-3 activation was determined by the translocation of mCardinal from cytoplasm into nucleus.

For imaging of the DEAD stained HeLa cells with PhoCl inserted Bid expression, photoconversion was performed under the 405 nm LED flood array with 15 s illumination. Following photoconversion, the cells were incubated in HHBSS buffer (with or without Z-VAD-FMK) at room temperature for 30 min. The working concentration of Z-VAD-FMK (Sigma-Aldrich) was 20 µM. For analysis the cell death, the DEAD Cytotoxicity kit for mammalian cells (Invitrogen) was used to stain the dead cells with or without photoconversion in HHBSS buffer (with or without Z-VAD-FMK) for 30 min at room temperature according to the manufacturer’s protocol. Images were acquired in white light, green fluorescence and red fluorescence channels. The extent of cell death was determined by the ratio of DEAD reagent stained cells to the green fluorescent transfected cells.

### Statistical analysis

All data are expressed as mean ± SEM. Sample sizes (*n*), significant differences (*P*) and confidence of curve fitting (*R*-squared) are listed for each experiment. Results are reported as *P* = *P* value, and *t* (degrees of freedom (df)) = *t* value. For all statistics, NS = *P* ≥ 0.05, * = *P* < 0.05, ** = *P* < 0.01, *** = *P* < 0.001 and **** = *P* < 0.0001. No samples were excluded from analysis and all experiments were reproducible. No randomization or blinding was used. The analyses were performed using GraphPad Prism for Mac (version 8.0.0), GraphPad Software, San Diego, California USA, www.graphpad.com.

## Data availability

Expression vectors for the following gene products and their gene sequences are available via Addgene and GenBank: pET-PhoCl1-6His, Addgene ID: 164033, and GenBank accession no. MW296870; pBad-LgBiT-PhoCl1-SmBiT-MBP, Addgene ID: 164034, and GenBank accession no. MW307773; pBad-PhoCl2c, Addgene ID: 164035, and GenBank accession no. MW296871; pBad-PhoCl2f, Addgene ID: 164036, and GenBank accession no. MW307774; pcDNA-NES-PhoCl2c-mCherry, Addgene ID: 164037, and GenBank accession no. MW307775; pcDNA-NES-PhoCl2f-mCherry, Addgene ID: 164048, and GenBank accession no. MW307776; pcDNA-NBid-mMaple-CBid, Addgene ID: 164049, and GenBank accession no. MW307777; pcDNA-NBid-PhoCl1-CBid, Addgene ID: 164050, and GenBank accession no. MW307778; pcDNA-NBid-PhoCl2c-CBid, Addgene ID: 164051, and GenBank accession no. MW307779; pcDNA-NES-DEVD-mCardinal-NLS, Addgene ID: 164052, and GenBank accession no. MW307780. Other expression vectors that support the findings in this study are available from the corresponding author on request. The primers used in this study and nucleotide sequences of all constructs and the are available in **Supplementary Table 7**,**8**. The coordinates of the PhoCl1 green, red and empty barrel structures are deposited in the Protein Data Bank under the codes of 7DMX, 7DNA and 7DNB, respectively.

## Acknowledgements

This work was supported by grants from the Canadian Institutes of Health Research (FS-154310 to REC) and the Natural Sciences and Engineering Research Council of Canada (RGPIN 2018-04364 to REC and RGPIN-2016-06478 to MJL). Y. W. is supported by the Alberta Parkinson Society Fellowship and National Natural Science Foundation of China (NO. 31870132 and NO. 82072237). S. Z. is supported by NSERC CREATE Advanced Protein Engineering Training, Internships, Courses, and Exhibition (APRENTICE) program. Part of the research described in this paper was performed using beamline CMCF-ID at the Canadian Light Source, a national research facility of the University of Saskatchewan, which is supported by the Canada Foundation for Innovation (CFI), the Natural Sciences and Engineering Research Council (NSERC), the National Research Council (NRC), the Canadian Institutes of Health Research (CIHR), the Government of Saskatchewan, and the University of Saskatchewan. This research was enabled in part by support provided by WestGrid (https://www.westgrid.ca/) and Compute Canada (www.computecanada.ca). We thank the University of Alberta Molecular Biology Services Unit, Gareth Lambkin, and Landon Zarowny for technical assistance. The furimazine luciferase substrate was synthesized by the Canadian Glycomics Network.

## Author Contributions

L. developed PhoCl2 variants, assembled all constructs, performed directed evolution, performed protein characterization, performed live cell imaging experiments, analyzed data, prepared figure, and wrote the manuscript. Y. W. performed the protein crystallization, solved the structures, and wrote the manuscript. S. Z. performed the molecular dynamic simulation, prepared figures, and wrote the manuscript. W. Z. conceived the idea of light-induced cell death experiment, performed the proof-of-concept tests, and prepared the figure. Y. C. performed the molecular dynamics simulation and the analysis. Y. S., M. J. L., and R. E. C. supervised research and edited the manuscript.

## References

1. Tischer, D. & Weiner, O. D. Illuminating cell signalling with optogenetic tools. Nat. Rev. Mol. Cell Biol. 15, 551–558 (2014).

2. Liu, Q. & Tucker, C. L. Engineering genetically-encoded tools for optogenetic control of protein activity. Curr. Opin. Chem. Biol. 40, 17–23 (2017).

3. Johnson, H. E. & Toettcher, J. E. Illuminating developmental biology with cellular optogenetics. Curr. Opin. Biotechnol. 52, 42–48 (2018).

4. Lu, X., Shen, Y. & Campbell, R. E. Engineering Photosensory Modules of Non-Opsin-Based Optogenetic Actuators. Int. J. Mol. Sci. 21, 6522 (2020).

5. Zhang, W. et al. Optogenetic control with a photocleavable protein, Phocl. Nat. Methods 14, 391–394 (2017).

6. Nagel, G. et al. Channelrhodopsin-2, a directly light-gated cation-selective membrane channel. Proc. Natl. Acad. Sci. U. S. A. 100, 13940–13945 (2003).

7. Gradinaru, V., Thompson, K. R. & Deisseroth, K. eNpHR: A Natronomonas halorhodopsin enhanced for optogenetic applications. Brain Cell Biol. 36, 129–139 (2008).

8. Gradinaru, V. et al. Molecular and Cellular Approaches for Diversifying and Extending Optogenetics. Cell 141, 154–165 (2010).

9. Airan, R. D., Thompson, K. R., Fenno, L. E., Bernstein, H. & Deisseroth, K. Temporally precise in vivo control of intracellular signalling. Nature 458, 1025–1029 (2009).

10. Niopek, D. et al. Engineering light-inducible nuclear localization signals for precise spatiotemporal control of protein dynamics in living cells. Nat. Commun. 5, (2014).

11. Niopek, D., Wehler, P., Roensch, J., Eils, R. & Di Ventura, B. Optogenetic control of nuclear protein export. Nat. Commun. 7, 1–9 (2016).

12. Wu, Y. I. et al. A genetically encoded photoactivatable Rac controls the motility of living cells. Nature 461, 104–108 (2009).

13. Strickland, D. et al. TULIPs: Tunable, light-controlled interacting protein tags for cell biology. Nat. Methods 9, 379–384 (2012).

14. Kawano, F., Suzuki, H., Furuya, A. & Sato, M. Engineered pairs of distinct photoswitches for optogenetic control of cellular proteins. Nat. Commun. 6, (2015).

15. Kennedy, M. J. et al. Rapid blue-light-mediated induction of protein interactions in living cells. Nat. Methods 7, 973–975 (2010).

16. Levskaya, A., Weiner, O. D., Lim, W. A. & Voigt, C. A. Spatiotemporal control of cell signalling using a light-switchable protein interaction. Nature 461, 997–1001 (2009).

17. Kaberniuk, A., Shemetov, A. A. & Verkhusha, V. V. A bacterial phytochrome-based optogenetic system controllable with near-infrared light. Nat. Methods 13, 591–597 (2016).

18. Zhou, X. X., Chung, H. K., Lam, A. J. & Lin, M. Z. Optical Control of Protein Activity by Fluorescent Protein Domains. Science. 338, 810–814 (2012).

19. Mizuno, H. et al. Photo-induced peptide cleavage in the green-to-red conversion of a fluorescent protein. Mol. Cell 12, 1051–1058 (2003).

20. Shadish, J. A., Strange, A. C. & Deforest, C. A. Genetically Encoded Photocleavable Linkers for Patterned Protein Release from Biomaterials. J. Am. Chem. Soc. 141, 15619– 15625 (2019).

21. Xiang, D. et al. Hydrogels With Tunable Mechanical Properties Based on Photocleavable Proteins. Front. Chem. 8, 1–9 (2020).

22. Endo, M., Iwawaki, T., Yoshimura, H. & Ozawa, T. Photocleavable Cadherin Inhibits Cell-to-Cell Mechanotransduction by Light. ACS Chem. Biol. 14, 2206–2214 (2019).

23. Reed, E. H., Schuster, B. S., Good, M. C. & Hammer, D. A. SPLIT: Stable Protein Coacervation Using a Light Induced Transition. ACS Synth. Biol. 9, 500–507 (2020).

24. Hall, M. P. et al. Engineered luciferase reporter from a deep sea shrimp utilizing a novel imidazopyrazinone substrate. ACS Chem. Biol. 7, 1848–1857 (2012).

25. Dixon, A. S. et al. NanoLuc Complementation Reporter Optimized for Accurate Measurement of Protein Interactions in Cells. ACS Chem. Biol. 11, 400–408 (2016).

26. Tsien, R. Y. The green fluorescent protein. Annu. Rev. Biochem. 67, 509–544 (1998).

27. Hori, Y. Crystal structure of the Aequorea victoria green fluorescent protein. Tanpakushitsu Kakusan Koso. 52, 1768–1769 (2007).

28. Nienhaus, K., Nienhaus, G. U., Wiedenmann, J. & Nar, H. Structural basis for photo-induced protein cleavage and green-to-red conversion of fluorescent protein EosFP. Proc. Natl. Acad. Sci. U. S. A. 102, 9156–9159 (2005).

29. Tsutsui, H. et al. The E1 Mechanism in Photo-Induced β-Elimination Reactions for Green-to-Red Conversion of Fluorescent Proteins. Chem. Biol. 16, 1140–1147 (2009).

30. McEvoy, A. L. et al. mMaple: A Photoconvertible Fluorescent Protein for Use in Multiple Imaging Modalities. PLoS One 7, (2012).

31. Xu, Y., Piston, D. W. & Johnson, C. H. A bioluminescence resonance energy transfer (BRET) system: Application to interacting circadian clock proteins. Proc. Natl. Acad. Sci. U. S. A. 96, 151–156 (1999).

32. Henderson, B. R. & Eleftheriou, A. A comparison of the activity, sequence specificity, and CRM1-dependence of different nuclear export signals. Exp. Cell Res. 256, 213–224 (2000).

33. Esposti, M. D. The roles of Bid. Apoptosis 7, 433–440 (2002).

34. Billen, L. P., Shamas-Din, A. & Andrews, D. W. Bid: A Bax-like BH3 protein. Oncogene 27, S93–S104 (2008).

35. Schug, Z. T., Gonzalvez, F., Houtkooper, R. H., Vaz, F. M. & Gottlieb, E. BID is cleaved by caspase-8 within a native complex on the mitochondrial membrane. Cell Death Differ. 18, 538–548 (2011).

36. Slee, E. A. et al. Benzyloxycarbonyl-Val-Ala-Asp (OMe) fluoromethylketone (Z-VAD.FMK) inhibits apoptosis by blocking the processing of CPP32. Biochem. J. 315, 21– 24 (1996).

37. Campbell, R. E. Fluorescent-Protein-Based Biosensors: Modulation of Energy Transfer as a Design Principle. Anal. Chem. 81, 5972–5979 (2009).

38. Wang, H. et al. LOVTRAP: An optogenetic system for photoinduced protein dissociation. Nat. Methods 13, 755–758 (2016).

39. Gross, L. A., Baird, G. S., Hoffman, R. C., Baldridge, K. K. & Tsien, R. Y. The structure of the chromophore within DsRed, a red fluorescent protein from coral. Proc. Natl. Acad. Sci. U. S. A. 97, 11990–11995 (2000).

40. Heim, R., Cubitt, A. B. & Tsien, R. Y. Improved green fluorescence. Nature 373, 663–664 (1995).

41. Kabsch, W. Integration, scaling, space-group assignment and post-refinement. Acta Crystallogr. Sect. D Biol. Crystallogr. 66, 133–144 (2010).

42. McCoy, A. J. Solving structures of protein complexes by molecular replacement with Phaser. Acta Crystallogr. Sect. D Biol. Crystallogr. 63, 32–41 (2006).

43. Ai, H. W., Henderson, J. N., Remington, S. J. & Campbell, R. E. Directed evolution of a monomeric, bright and photostable version of Clavularia cyan fluorescent protein: Structural characterization and applications in fluorescence imaging. Biochem. J. 400, 531–540 (2006).

44. Emsley, P. & Cowtan, K. Coot: Model-building tools for molecular graphics. Acta Crystallogr. Sect. D Biol. Crystallogr. 60, 2126–2132 (2004).

45. Adams, P. D. et al. PHENIX: A comprehensive Python-based system for macromolecular structure solution. Acta Crystallogr. Sect. D Biol. Crystallogr. 66, 213–221 (2010).

46. Strong, M. et al. Toward the structural genomics of complexes: Crystal structure of a PE/PPE protein complex from Mycobacterium tuberculosis. Proc. Natl. Acad. Sci. U. S. A. 103, 8060–8065 (2006).

47. Schrodinger LLC. The PyMOL Molecular Graphics System, Version 1.8. (2015).

48. Pettersen, E. F. et al. UCSF Chimera - A visualization system for exploratory research and analysis. J. Comput. Chem. 25, 1605–1612 (2004).

49. Wallace, A. C., Laskowski, R. A. & Thornton, J. M. LIGPLOT: a program to generate schematic diagrams of protein-ligand interactions. Protein Eng. 8, 127–134 (1995).

50. Hanwell, M. D. et al. Avogadro: an advanced semantic chemical editor, visualization, and analysis platform. J. Cheminform. 4, 17 (2012).

51. Vanquelef, E. et al. R.E.D. Server: A web service for deriving RESP and ESP charges and building force field libraries for new molecules and molecular fragments. Nucleic Acids Res. 39, 511–517 (2011).

52. Wink, L. H., Baker, D. L., Cole, J. A. & Parrill, A. L. A benchmark study of loop modeling methods applied to G protein-coupled receptors. J. Comput. Aided. Mol. Des. 33, 573–595 (2019).

53. Jarzynski, C. Nonequilibrium Equality for Free Energy Differences. Phys. Rev. Lett. 78, 2690–2693 (1997).

54. Roe, D. R. & Cheatham, T. E. PTRAJ and CPPTRAJ: Software for processing and analysis of molecular dynamics trajectory data. J. Chem. Theory Comput. 9, 3084–3095 (2013).

55. Miller, B. R. et al. MMPBSA.py: An efficient program for end-state free energy calculations. J. Chem. Theory Comput. 8, 3314–3321 (2012).

56. Humphrey, W., Dalke, A. & Schulten, K. VMD - Visual Molecular Dynamics. J. Mol. Graph. 14, 33–38 (1996).

57. Kalderon, D., Roberts, B. L., Richardson, W. D. & Smith, A. E. A short amino acid sequence able to specify nuclear location. Cell 39, 499–509 (1984).

