## Supplementary information for "Improved Photocleavable Proteins with Faster and More Efficient Dissociation"

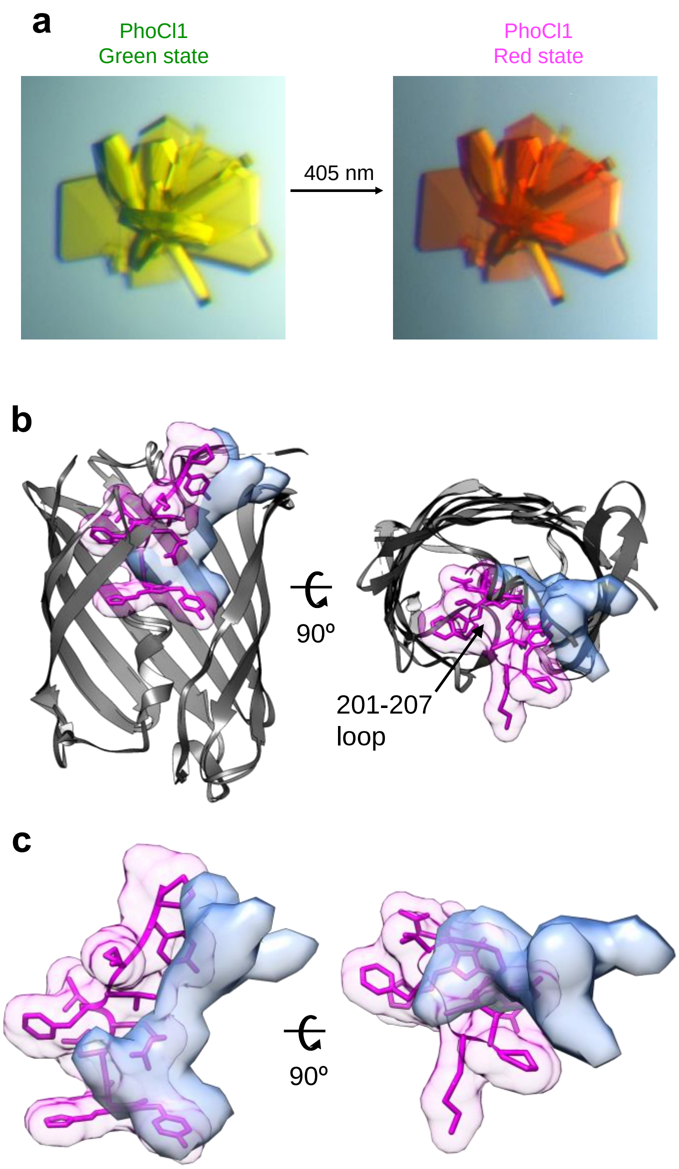


**Supplementary Figure 1**. Additional information on crystallization and conformational changes in the 201-207 loop. (**a**) Visible color changes in a crystal of PhoCl1 before and after illumination. Illumination with violet light (15 s light with LED array, 3.46 mW/mm$^2$). (**b**) Alternative view of the superimposed PhoCl1 red state and empty barrel. The β-barrel of the red state structure is represented in grey, with the dissociable peptide and chromophore shown as magenta sticks surrounded by a semi-transparent magenta surface. The empty barrel structure is represented in grey with the water-filled cavity represented as semi-transparent light blue surface. (**c**) Zoomed-in representation of the superimposed dissociable peptide and chromophore (semi-transparent magenta surface) in the red state and water-filled cavity (semi-transparent light blue surface) in the empty barrel.

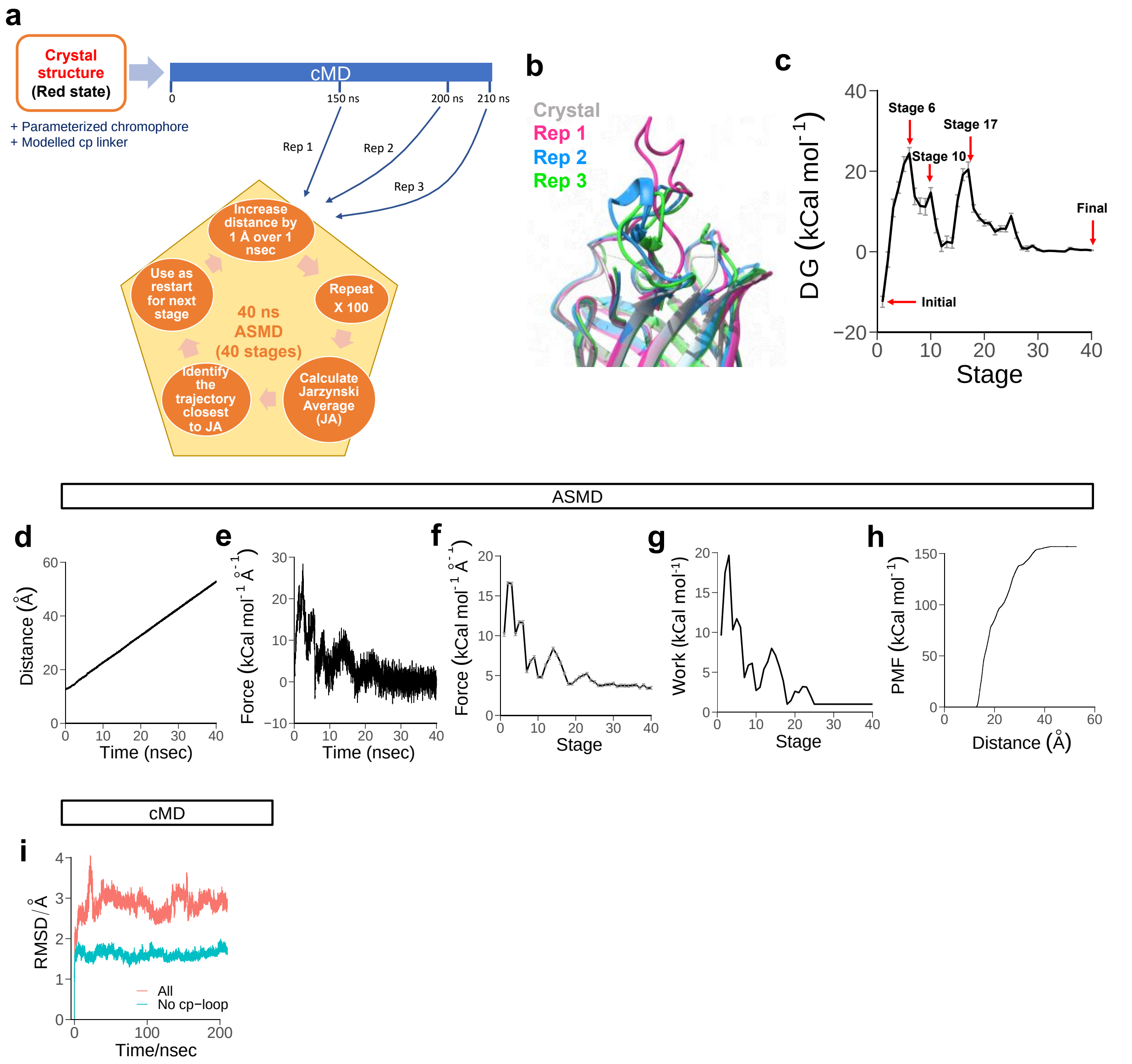


**Supplementary Figure 2**. Additional information on molecular dynamic simulation study of dissociation process. (**a**) Schematic of the workflow of molecular dynamic simulations. Rep, replication. (**b**) Structure alignment of the crystal structure of the PhoCl1 red state and the simulated initial stages of 3 ASMD replications with different conformations of the cp linker. The crystal structure of the PhoCl1 red state is shown in silver, Rep1 is shown in magenta, Rep2 is shown in blue and Rep3 is shown in green. (**c**) Gibbs free energy of activation over stages from Rep3 with the final stage as reference (ΔG = 0 kCal/mol). Values are means ± SEM ($n$ = 20 snapshots per stage). The stages represented in **Fig. 2a,b** are indicated by red arrows. (**d-h**) During the ASMD, (**d**) The distance between the COMs of the C-terminal peptide and the N-terminal barrel increased by 40 Å over 40 ns imposed by a harmonic restraint. With a constant speed of movement, the force imposing such distance constraint is changing. The curve is smoothened by (**e**) averaging the adjacent 10 snapshots or (**f**) averaging all snapshots within a stage for clarity. The work done by such force in each stage is shown in (**g**). The cumulative potential of mean force (PMF) throughout the ASMD is shown in (**h**). (**i**) During the cMD preceeding the ASMD, the conformation dynamics of the structure is shown with the RMSD of all residues with or without the cp linker. The analysis suggests that most of the movements take place at the cp linker while the rest of the protein remains largely still in cMD, including the 201-207 loop.

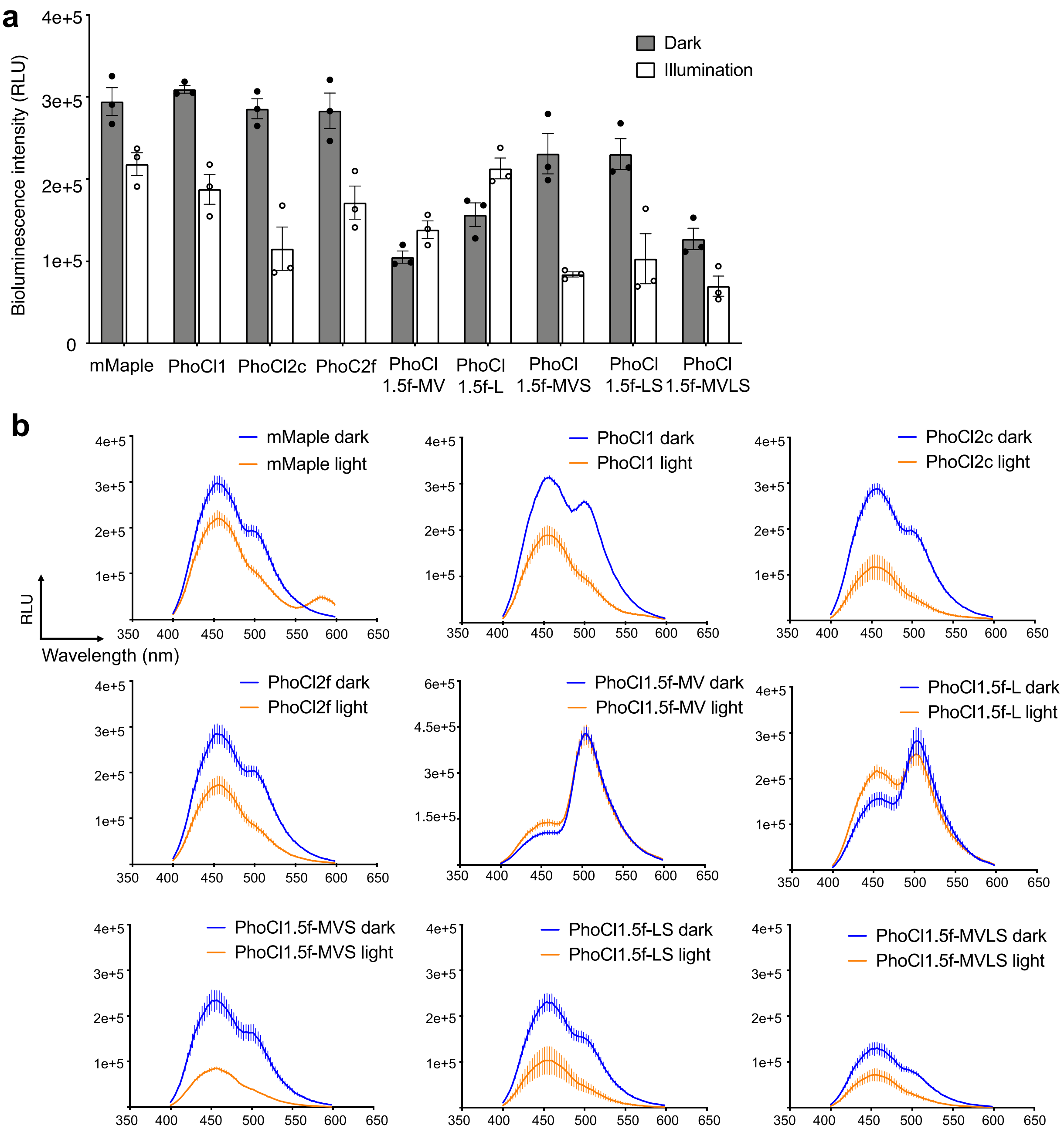


**Supplementary Figure 3**. Bioluminescence assay of PhoCl inserted NanoBiT. (**a**) Summary data of bioluminescence intensity (at 460 nm) on key variants. Purified LgBiT-PhoCl-SmBiT-MBP fusion protein (500 nM; construct represented in **Fig. 3a**) was applied in this assay. At 30 min after illumination (15 s with LED array), the luminescence emission spectra of the protein, both with and without photoconversion, were measured immediately after treatment with the luciferase substrate furimazine (final concentration: 10 µg/mL). Values are means ± SEM (*n* = 3 independent experiments). (**b**) Bioluminescence spectra of PhoCl inserted NanoBiT with (light) or without (dark) illumination. Measurement details were described in **a**.

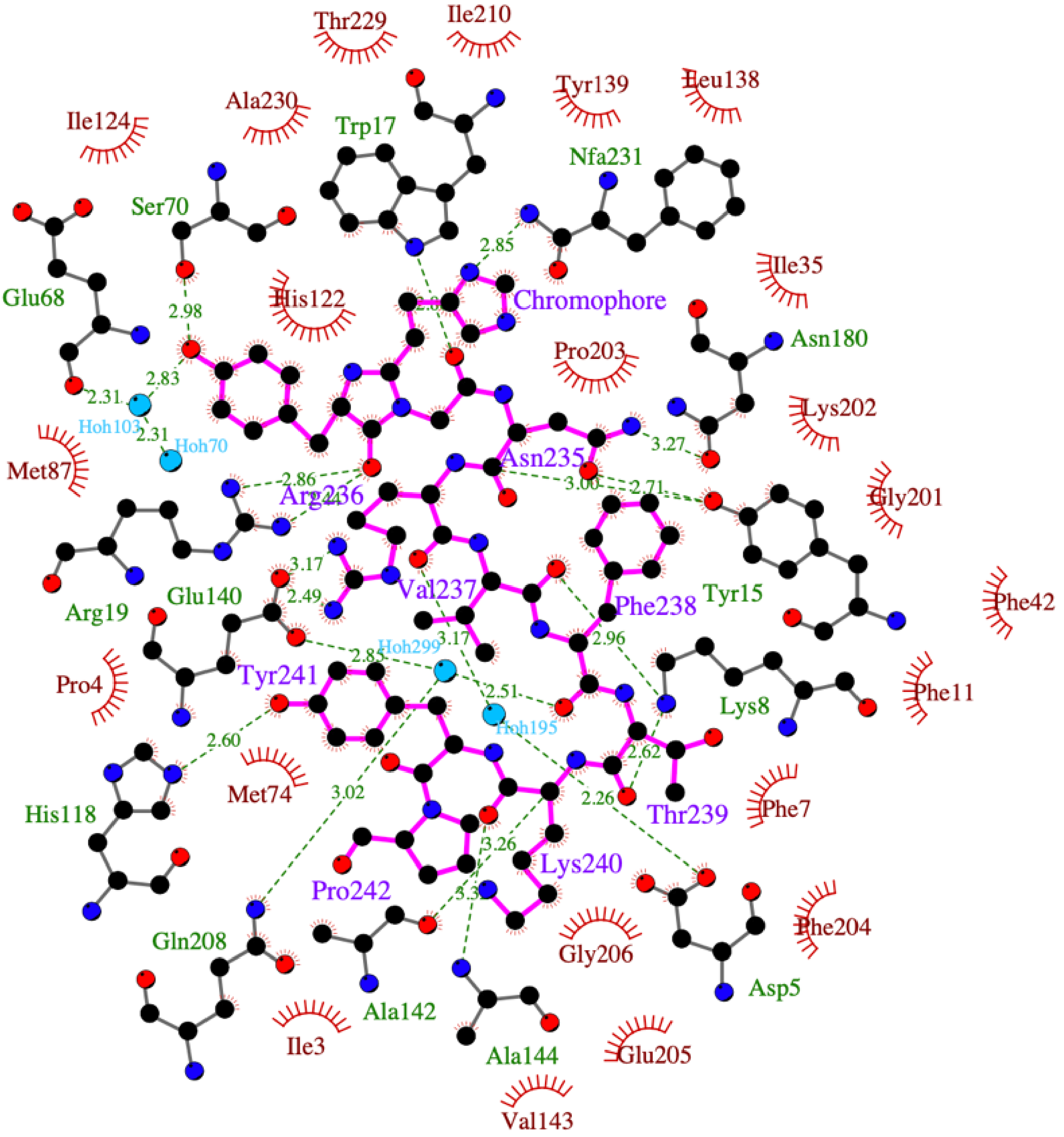


**Supplementary Figure 4**. Schematic diagram of the dissociable peptide and chromophore in the PhoCl1 red state generated by LIGPLOT[1]. The peptide and chromophore are represented with magenta-colored bonds. Hydrogen bonds are represented in green dashed lines and the hydrogen-bonded residues are represented with grey-colored bonds. Hydrophobic contacts are represented as red arcs with radiating lines.

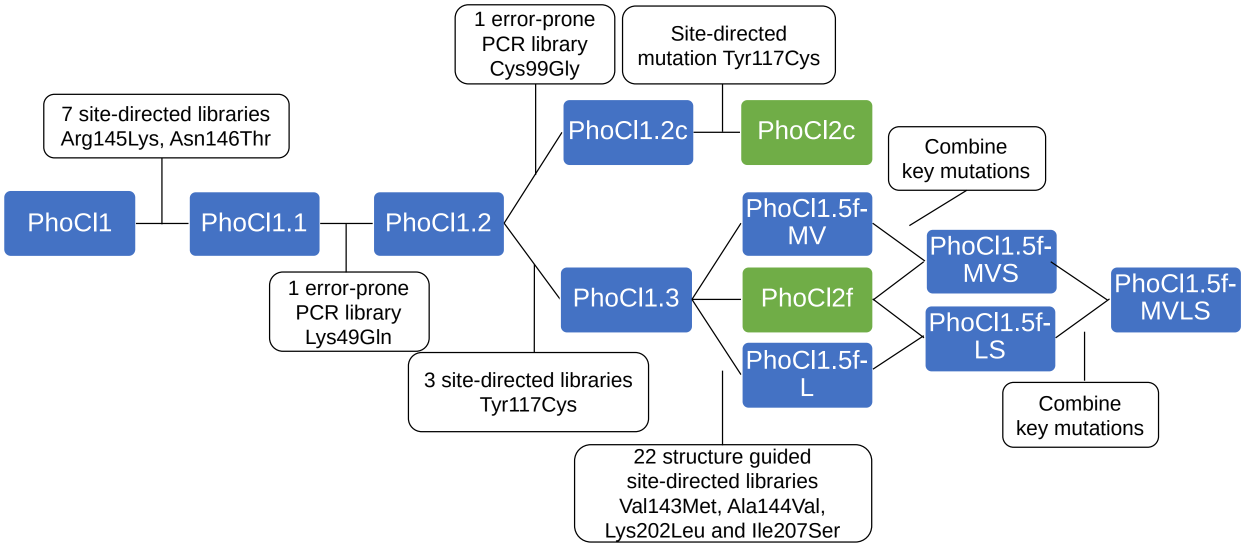


**Supplementary Figure 5**. Flow chart of PhoCl evolution. The PhoCl variants in the screening process are represented in blue rectangles. The final PhoCl2 variants are presented in green rectangles. The library generation method, specific mutations discovered, and other key points, are represented in white rectangles.

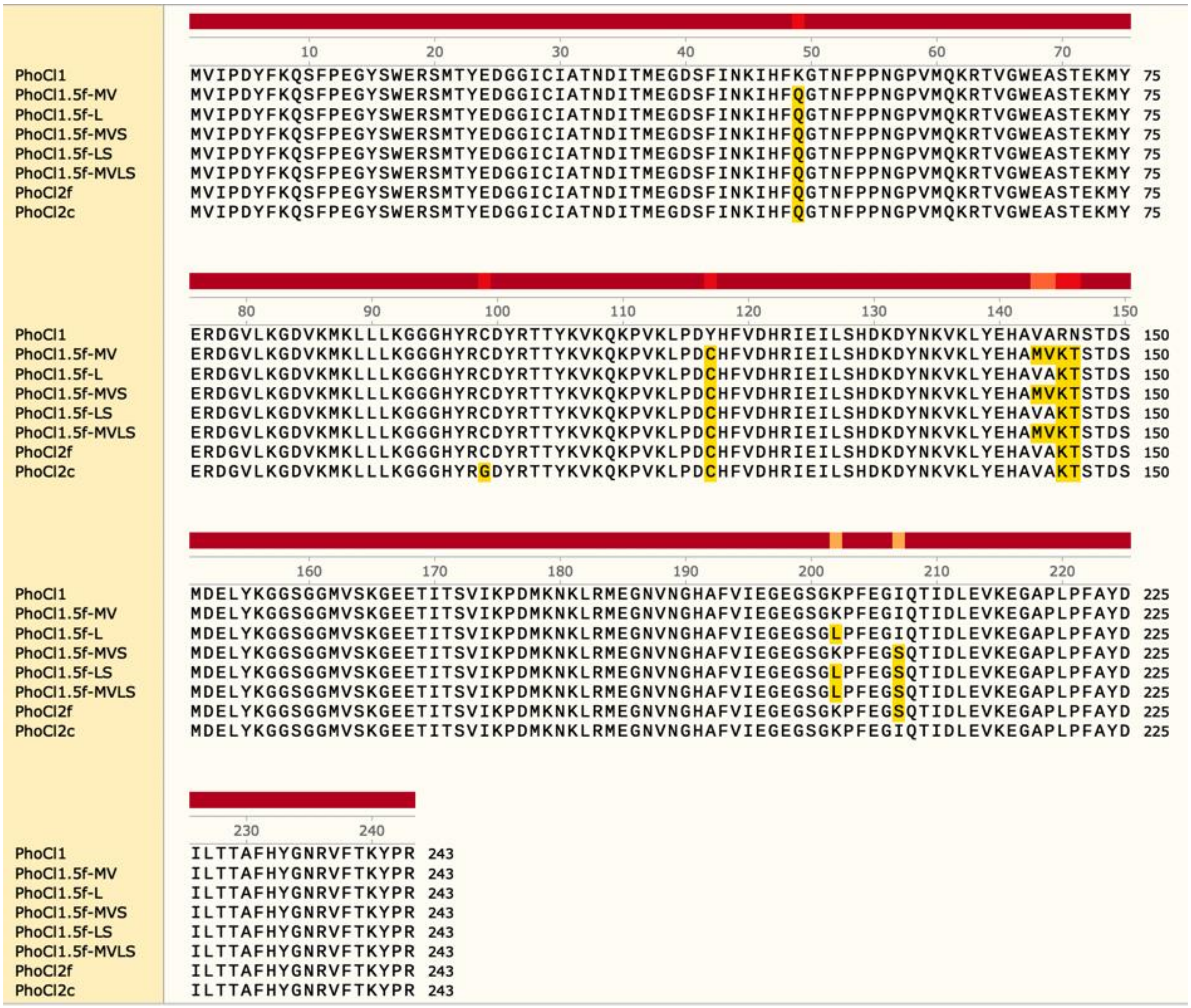


**Supplementary Figure 6**. Sequence alignment of PhoCl variants. Alignment was performed on SnapGene software. The sequence of PhoCl1 was used as reference. The mutations in variants were highlighted in yellow-orange.

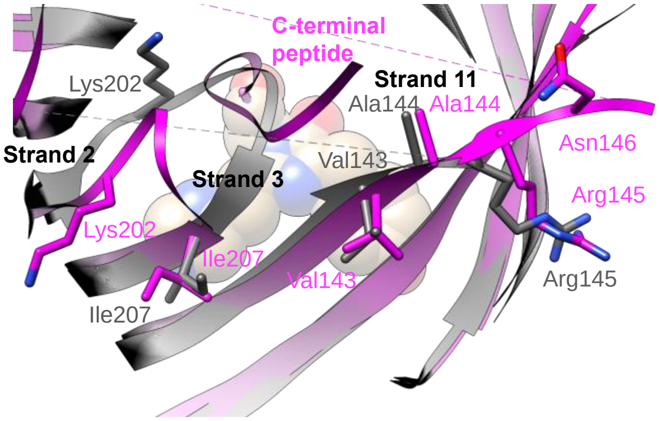


**Supplementary Figure 7**. Key mutations in or near the 201-207 loop that were identified during screening.

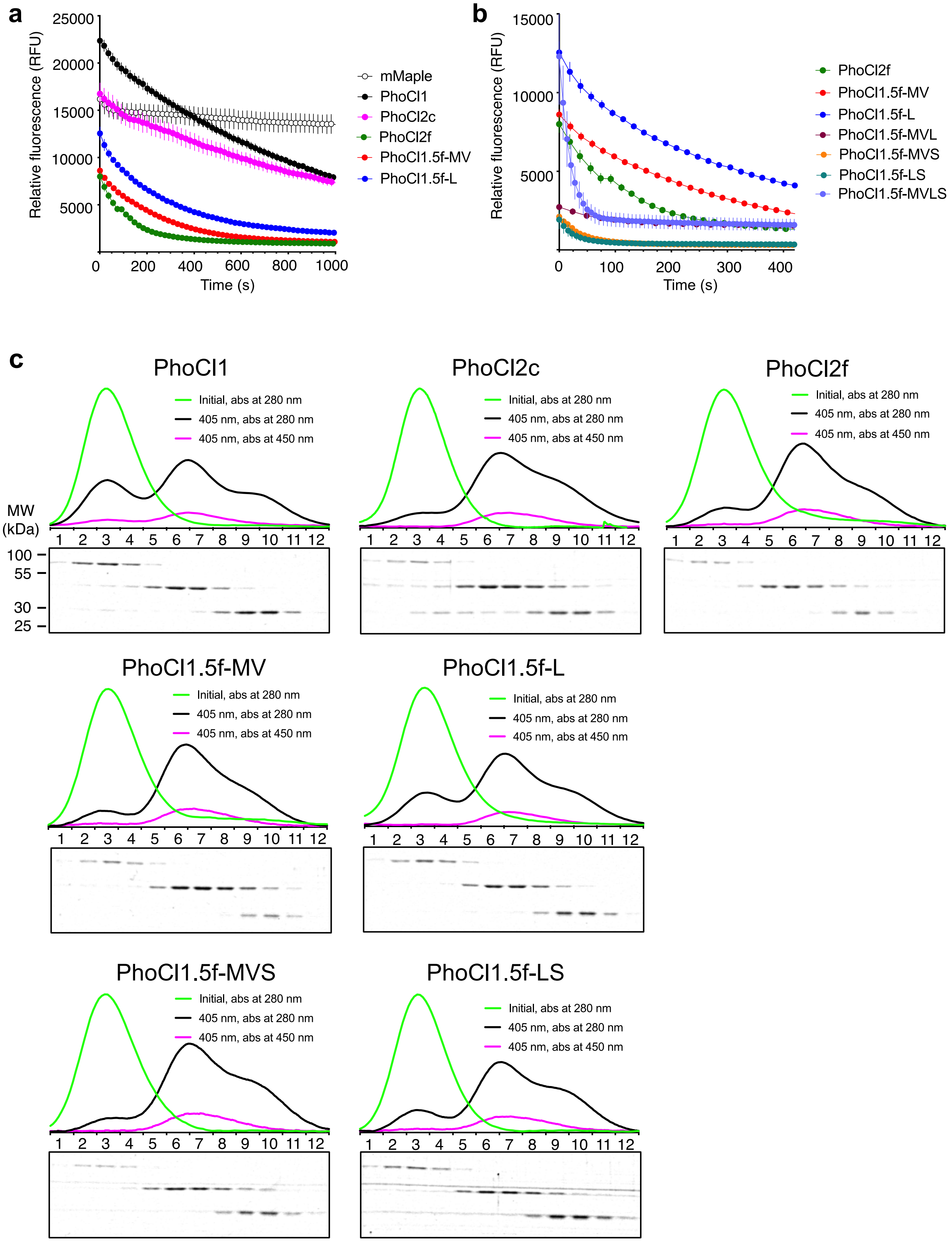


**Supplementary Figure 8**. Additional data on dissociation kinetics and efficiencies of PhoCl variants. **(a,b)** Loss of red fluorescence after photoconversion without normalization. This is the same data shown in **Fig. 3b** (panel a) and **Fig. 3c** (panel b). RFU, relative fluorescence units. Each protein was at a concentration of 500 nM, except for PhoCl1.5f-MVLS which was at 5 µM due to its poor chromophore formation. **(c)** GFC and SDS-PAGE analysis of PhoCl-MBP fusions. SDS-PAGE analysis of GFC fractions for partial photoconverted fusion protein is labeled by fraction numbers (12 × 1.5 mL, 43.5 - 61.5 mL elution volume).

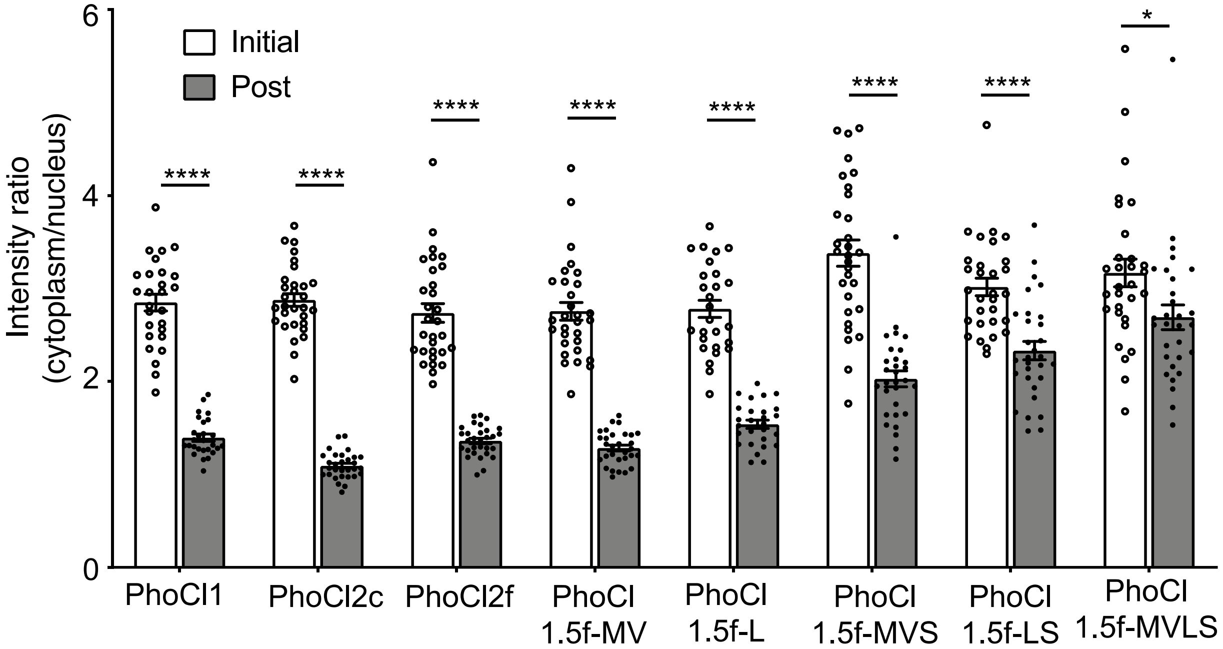


**Supplementary Figure 9**. Red fluorescence intensity localization ratios (cytoplasm to nucleus) of NES-PhoCl-mCherry before and after photoconversion. Values are means ± SEM. The data is from the same experiments shown in **Fig. 4**. PhoCl1: $****P < 0.000001$, $t\ (52) = 14.85$, $n = 27$ cells from 3 cultures; PhoCl2c: $****P < 0.000001$, $t\ (58) = 24.41$, $n = 30$ cells from 3 cultures; PhoCl2f: $****P < 0.000001$, $t\ (58) = 13.09$, $n = 30$ cells from 3 cultures; PhoCl1.5f-MV: $****P < 0.000001$, $t\ (58) = 14.47$, $n = 30$ cells from 3 cultures; PhoCl1.5f-L: $****P < 0.000001$, $t\ (52) = 12.03$, $n = 27$ cells from 3 cultures; PhoCl1.5f-MVS: $****P < 0.000001$, $t\ (58) = 8.187$, $n = 30$ cells from 3 cultures; PhoCl1.5f-LS: $****P = 0.000006$, $t\ (58) = 4.997$, $n = 30$ cells from 3 cultures; PhoCl1.5f-MVLS: $*P = 0.020512$, $t\ (58) = 2.382$, $n = 30$ cells from 3 cultures. Multiple *t* tests were used to analyze significant difference between group means.

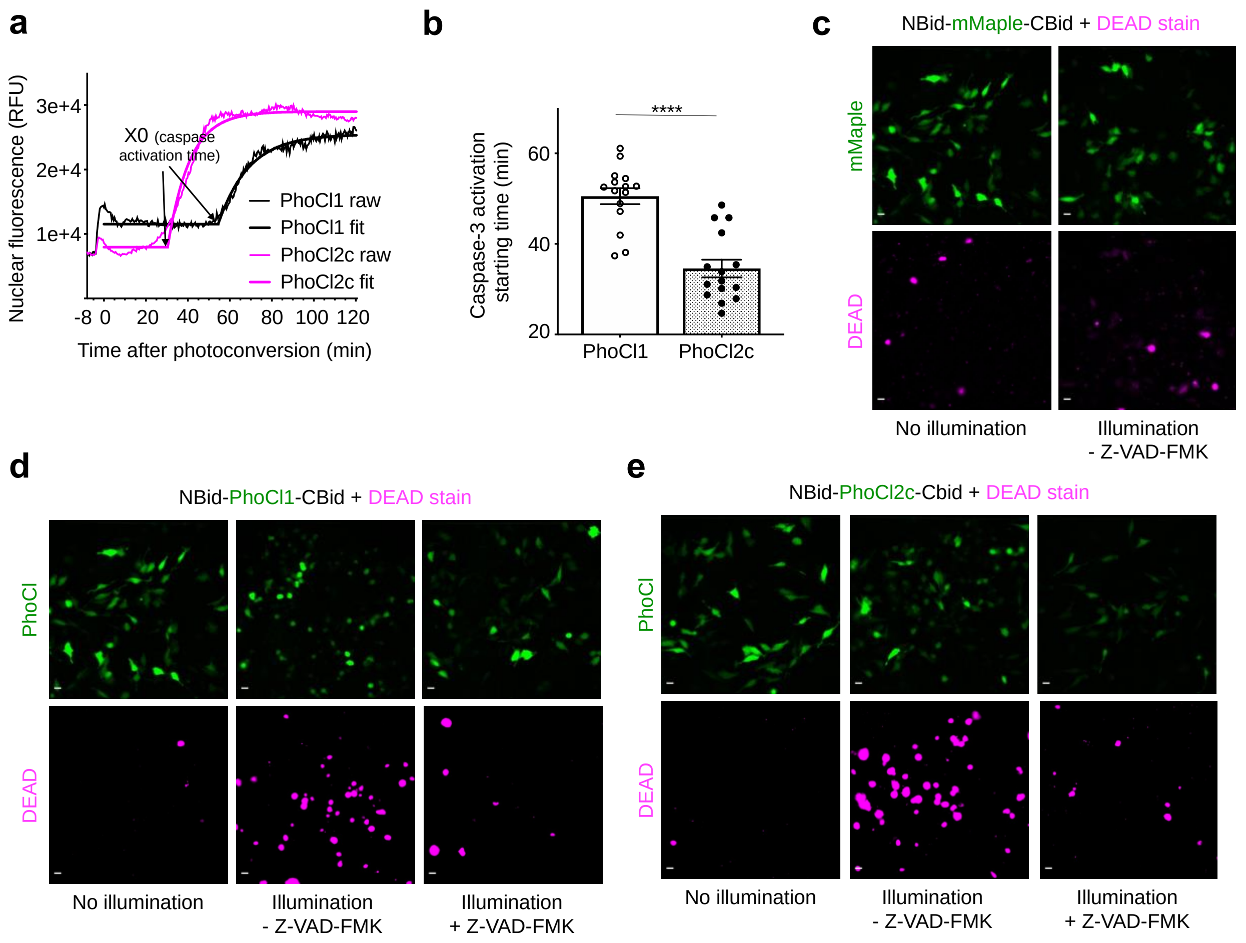


**Supplementary Figure 10**. Demonstration of PhoCl-dependent induction of apoptosis using the caspase-3 translocation reporter and DEAD cell viability assay. (**a**) Representative nuclear fluorescence intensity profiles of the HeLa cell co-expressing PhoCl inserted Bid and the NES-DEVD-mCardinal-NLS caspase-3 reporter represented in **Fig. 5c**. RFU, relative fluorescence units. Cells were illuminated with 10 s violet light pulses (395/40 nm, 2 mW/mm$^2$) every 15 s for 6 mins, then imaged 2 hours after photoconversion. Plateau followed by one phase decay fit was applied to each nuclear fluorescence profile. $R$-squared values range from 0.8238 to 0.9940. The caspase-3 activation time was characterized by X0 in the fit. The activation time X0 in each nuclear fluorescence profile was plotted in **b**. (**b**) Summary of caspase-3 activation starting time for cells in **Fig. 5c**. Values are mean ± SEM ($n$ = 15 cells from 3 cultures for each variant). ****$P$ < 0.0001 by unpaired two-tailed *t* test (*t* (28) = 6.043). (**c-e**) Representative cell images of HeLa cells expressing mMaple or PhoCl inserted Bid after DEAD stain. Scale bar, 20 μm.

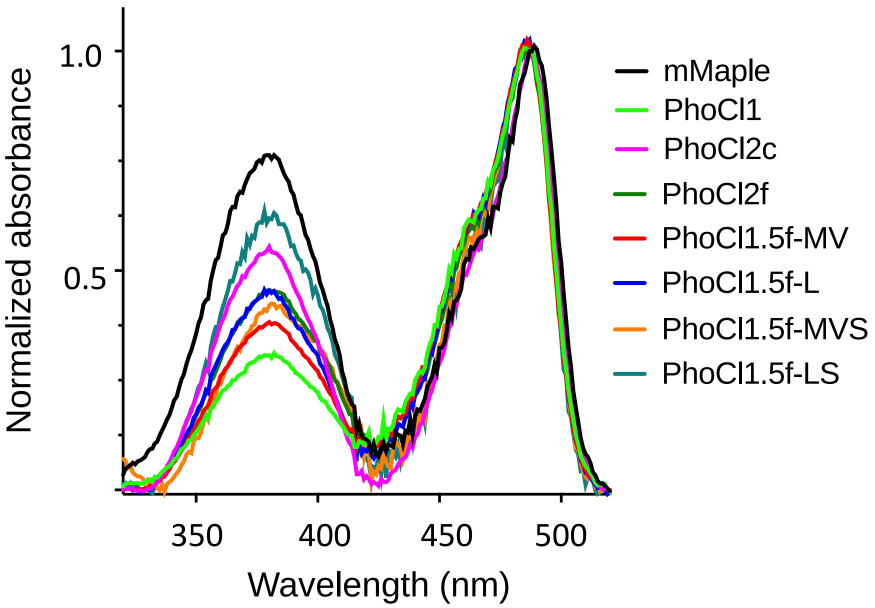


**Supplementary Figure 11**. Absorbance spectra of mMaple and PhoCl variants. Compared to PhoCl1, increased extinction coefficients at 405 nm were observed with the improved variants.

**Supplementary Table 1.** X-ray data collection and refinement statistics.

| **Crystal** | **PhoCl1 green state** | **PhoCl1 red state** | **PhoCl1 empty barrel** |
|---|---|---|---|
| **Data collection** | | | |
| Spacegroup | $P2_12_12_1$ | P1 | $P2_1$ |
| a, b, c (Å) | 60.36, 112.76, 144.85 | 38.93, 72.49, 126.55 | 46.4, 119.6, 65.3 |
| α, β, γ (°) | 90.0, 90.0, 90 | 92.5, 97.3, 92.5 | 90, 107.7, 90 |
| Resolution (Å) | 42.88-2.10 (2.18-2.10) | 38.57-2.30 (2.38-2.30) | 43.11-2.82 (2.92-2.82) |
| *R*merge | 0.066 (0.709) | 0.042 (0.533) | 0.073 (0.760) |
| *R*meas | 0.078 (0.835) | 0.059 (0.734) | 0.089 (0.936) |
| Multiplicity | 3.4 (3.5) | 1.8 (1.7) | 2.7 (2.7) |
| CC(1/2) | 0.997 (0.885) | 0.998 (0.855) | 0.997 (0.811) |
| CC* | 0.999 (0.969) | 1 (0.96) | 0.999 (0.946) |
| I/σ(I) | 9.73 (1.72) | 9.60 (1.20) | 8.54 (1.28) |
| Completeness (%) | 96.8 (97.7) | 95.9 (94.9) | 92.9 (94.5) |
| Wilson B-factor ($Å^2$) | 25.64 | 28.86 | 43.91 |
| **Refinement** | | | |
| Total Reflections | 194926 (31020) | 103548 (16288) | 44693 (7133) |
| Unique Reflections | 57043 (9153) | 58440 (9388) | 16400 (2688) |
| $R_{Work}/R_{Free}$ | 0.1824/0.2162 | 0.2186/0.2638 | 0.2587/0.3029 |
| **Number of atoms:** | | | |
| Protein | 6715 | 10075 | 5031 |
| Ligands | 100 | 148 | - |
| Water | 410 | 488 | - |
| Average B-factor ($Å^2$) | 34.07 | 35.53 | 38.70 |
| Protein ADP ($Å^2$) | 34.35 | 35.59 | 38.71 |
| Ligands ($Å^2$) | 22.13 | 29.24 | - |
| Water | 32.51 | 31.77 | - |
| **Ramachandran plot:** | | | |
| Favored/Allowed (%) | 97.70/2.20 | 94.0/5.4 | 91.0/8.7 |
| **Root-Mean-Square-Deviation:** | | | |
| Bond lengths (Å) | 0.012 | 0.004 | 0.005 |
| Bond Angle (°) | 1.50 | 1.0 | 0.82 |

*Note:* Statistics for the highest resolution shell are shown in parentheses.

**Supplementary Table 2**. Summary of the interactions of PhoCl1 dissociable peptide and chromophore in the red state.

| **Residues** | **Hydrogen Bonding** | **Hydrophobic Interaction (2.90 Å – 3.90 Å)** |
|---|---|---|
| Chromophore | Trp17 (2.86), Arg19 (2.86, 2.44), Ser70 (2.98), Nfa231 (2.85) | Met87, His122, Ile124, Leu138, Tyr139, Glu140, Gln208, Ile210, Thr229, Ala230, Arg236 |
| Asn235 | Tyr15 (2.71), Asn180 (3.27), Val237 (3.30) | Ile35 |
| Arg236 | Tyr15 (3.00), Glu140 (3.17, 2.49), Tyr239 (2.92,2.68) | Phe7, Met74, His122, Ala142, Iey233, Phe238 |
| Val237 | Lys8 (2.96), Asn235 (3.30) | Phe11, Phe42, Pro203, Thr239 |
| Phe238 | NA | Ala142, Asn180, Gly201, Lys202, Pro203, Gly206, Gln208, Arg235 |
| Thr239 | Lys8 (2.62), Arg236 (2.68, 2.92), Tyr241 (3.12) | Pro203, Gly206, Val237 |
| Lys240 | Ala142 (3.26), Ala144 (3.32) | Val143, Phe204, Glu205, Gly206 |
| Tyr241 | His118 (2.60), Thr239 (3.12) | Ile3, Asp5, Pro4 |
| Pro242 | NA | NA |

**Supplementary Table 3**. Summary table of mutations in PhoCl2 variants.

| Variants | Mutations |
|---|---|
| PhoCl1.5f-MV | Lys49Gln, Tyr117Cys, Val143Met, Ala144Val, Arg145Lys, Asn146Thr |
| PhoCl1.5f-L | Lys49Gln, Tyr117Cys, Arg145Lys, Asn146Thr, Lys202Leu |
| PhoCl1.5f-MVS | Lys49Gln, Tyr117Cys, Val143Met, Ala144Val, Arg145Lys, Asn146Thr, Ile207Ser |
| PhoCl1.5f-LS | Lys49Gln, Tyr117Cys, Arg145Lys, Asn146Thr, Lys202Leu, Ile207Ser |
| PhoCl1.5f-MVLS | Lys49Gln, Tyr117Cys, Val143Met, Ala144Val, Arg145Lys, Asn146Thr, Lys202Leu, Ile207Ser |
| PhoCl2f | Lys49Gln, Tyr117Cys, Arg145Lys, Asn146Thr, Ile207Ser |
| PhoCl2c | Lys49Gln, Cys99Gly, Tyr117Cys, Arg145Lys, Asn146Thr |

**Supplementary Table 4**. Properties of PhoCl variants.

| Protein | Extinction Coefficients ε (488 nm) | ε (405 nm) | Quantum Yield 488 nm | Brightness 488 nm | Dissociation Half Time (s) |
|---|---|---|---|---|---|
| mMaple | 22 | 9.3 | 0.89 | 20 | NA |
| PhoCl1 | 34 | 8.3 | 0.73 | 25 | 570 |
| PhoCl2c | 22 | 9.9 | 0.48 | 10 | 700 |
| PhoCl2f | 24 | 13 | 0.50 | 12 | 76 |
| PhoCl1.5f-MV | 27 | 11 | 0.53 | 15 | 130 |
| PhoCl1.5f-L | 25 | 12 | 0.45 | 11 | 140 |
| PhoCl1.5f-MVS | 25 | 13 | 0.29 | 7.3 | 30 |
| PhoCl1.5f-LS | 20 | 15 | 0.36 | 7.2 | 21 |
| PhoCl1.5f-MVLS | NA | NA | NA | NA | 14 |

*Note:* Extinction coefficient and brightness in $mM^{-1}cm^{-1}$. All measurements were performed with purified proteins in PBS solution (pH 7.4). EGFP extinction coefficient of 56 $mM^{-1}cm^{-1}$ (Ref. 2) and quantum yield of 0.6 (Ref. 3) at 488 nm were used as reference standards.

**Supplementary Table 5**. Summary data of bioluminescence assay.

| Protein | Emission (460 nm) | BRET ratio (dark) | BRET ratio (light) |
|---|---|---|---|
| mMaple | -26% | 0.63 | 0.44 |
| PhoCl1 | -39% | 0.82 | 0.47 |
| PhoCl2c | -60% | 0.66 | 0.40 |
| PhoCl2f | -39% | 0.69 | 0.45 |
| PhoCl1.5f-MV | +32% | 4.08 | 3.07 |
| PhoCl1.5f-L | +36% | 1.80 | 1.18 |
| PhoCl1.5f-MVS | -64% | 0.68 | 0.43 |
| PhoCl1.5f-LS | -55% | 0.63 | 0.41 |
| PhoCl1.5f-MVLS | -45% | 0.58 | 0.38 |

*Note:* The BRET ratio was determined by the use of the PhoCl (505 nm) to NanoBiT (460 nm) ratio (acceptor/donor ratio). All the data were the means from triplicate experiments for each variant.

**Supplementary Table 6**. Summary data of optogenetic manipulation of protein translocation assay with PhoCl variants.

| Protein | Photoconversion Half Time (s) | Dissociation Half Time (s) |
|---|---|---|
| PhoCl1 | 75.5 | 241 |
| PhoCl2c | 62.7 | 114 |
| PhoCl2f | 78.1 | 135 |
| PhoCl1.5f-MV | 83.6 | 160 |
| PhoCl1.5f-L | 69.6 | 155 |

Plateau followed by one phase decay fit was used in both analyses. For fit of photoconversion, *R*-squared values range from 0.9900 to 0.9951. For fit of dissociation, *R*-squared values range from 0.7006 to 0.9040.

**Supplementary Table 7**. Primers used in this study.

| Primers | Sequence 5'-3' |
|---|---|
| **pBAD/HisB-LgBiT-PhoCl-SmBiT-MBP construct** | |
| F-XhoI-LgBiT | GCAGGCTCGAGGATGGTCTTCACACTCGAAGATTTCGTTGGG |
| R-LgBiT-linker1-KpnI-PhoCl overlap PCR | GAAGTAGTCAGGGATCACGGTACCAGGTTCTCCGCCGCTAGAACCGCCTCCGCTACTCCC |
| F-linker1-Kpn1-PhoCl overlap PCR | GGCGGAGAACCTGGTACCGTGATCCCTGACTACTTCAAGCAGAGC |
| R1-PhoCl-XbaI-linker2-SmBiT | CACGCCGCCACTGCTTCCGCCACCGGACGAACCTCTAGACCGTGGGTACTTGGTGAACAC |
| R2-linker2-SmBiT-EcoRI | CAGCTGAATTCCAGGATCTCTTCAAAAAGTCTGTATCCAGTCACGCCGCCACTGCTT |
| F-EcoRI-linker3-MBP | GTACGGAATTCGGGGGTGGAGGTTCAAAAATCGAAGAAGGTAAACTGGTAATCTGGAT |
| R-MBP*-HindIII | CAGCCAAGCTTTTAAGTCTGCGCGTCTTTCAGGG |
| **Error-prone PCR for PhoCl libraries** | |
| EP-F-KpnI-PhoCl | CGGAGAACCTGGTACCGTGATCCCT |
| EP-R-XbaI-PhoCl | CGGACGAACCTCTAGACCGTGGGTA |
| **QuickChange primers for PhoCl libraries** | |
| I3-antisense | TCTGCTTGAAGTAGTCAGGMNNCACGGTACCAGGTTCTCCG |
| D5-sense | GAACCTGGTACCGTGATCCCTNNKTACTTCAAGCAGAGCTTCCCC |
| F7-antisense | CCTCGGGGAAGCTCTGCTTMNNGTAGTCAGGGATCACGGTA |
| K8-antisense | GCCCTCGGGGAAGCTCTGMNNGAAGTAGTCAGGGATCAC |
| F11-antisense | CCAGCTGTAGCCCTCGGGMNNGCTCTGCTTGAAGTAGTC |
| G14 Y15-antisense | AGGTCATGCTGCGCTCCCAGCTMNNMNNCTCGGGGAAGCTCTGCTTGAAG |
| I35-antisense | CTGTCCCCCTCCATTGTMNNGTCGTTGGTGGCGATGC |
| M37 E38-antisense | TGATGAAGCTGTCCCCMNNMNNTGTGATGTCGTTGGTGGCGATGCAGATGCCG |
| G39 D40-antisense | CTGGAAGTGGATCTTGTTGATGAAGCTMNNMNNCTCCATTGTGATGTCGTTGGTGGCGAT |
| F42-antisense | CTTGAAGTGGATCTGGTTGATMNNGCTGTCCCCCTCCATTGTGAT |
| R77 D78-antisense | TCACGTCGCCCTTCAGCACGCCMNNMNNCTCGTACATCTTCTCGGTGCTG |
| G79 V80-antisense | CTTCATCTTCACGTCGCCCTTCAGMNNMNNGTCGCGCTCGTACATCTTCTCGGT |
| M87-sense | GCCCTTCAGCAGCAGCTTMNNCTTCACGTCGCCCTTCAG |
| Y117-antisense | CGGTGGTCCACGAAGTGMNNGTCGGGCAGCTTTACGG |
| Y117 H118-antisense | TCTCGATGCGGTGGTCCACGAAMNNMNNGTCGGGCAGCTTTACGGGCTTC |
| V120-antisense | TCTCGATGCGGTGGTCMNNGAAGTGGCAGTCGGGC |
| H122-antisense | GCTCAGGATCTCGATGCGMNNGTCCACGAAGTGGCAGTC |
| L138 Y139-antisense | GGAAGTCTTGGCCACGGCGTGCTCMNNMNNCTTCACCTTGTTGTAGTCCTTGTC |
| V143 A144-antisense | GTCCATGCTGTCGGTGGAAGTCTTMNNMNNGGCGTGCTCGTACAGCTTCACCTT |
| R145 N146-antisense | CTCGTCCATGCTGTCGGTGGAMNNMNNGGCCACGGCGTGCTCGTACAG |
| M162-sense | GGTGGCAGCGGTGGCNNKGTGAGCAAGGGCGAG |
| D177 M178-sense | GAGACCATTACAAGCGTGATCAAGCCTNNKNNKAAGAACAAGCTGCGCATGGAGGGCAAC |
| G201 K202-antisense | CGTCTGGATGCCCTCGAAGGGMNNMNNGCTGCCCTCGCCCTCGATCAC |
| P203 F204-antisense | ATCAATCGTCTGGATGCCCTCMNNMNNCTTGCCGCTGCCCTCGCCCTC |
| F204-antisense | CGTCTGGATGCCCTCMNNGGGCTTGCCGCTGCC |
| E205 G206-antisense | ACCTCCAAATCAATCGTCTGGATMNNMNNGAAGGGCTTGCCGCTGCCCTCGC |
| I207-antisense | CTCCAAATCAATCGTCTGMNNGCCCTCGAAGGGCTTGCC |
| I207 Q208-antisense | TCCTTCACCTCCAAATCAATCGTMNNMNNGCCCTCGAAGGGCTTGCCGCTGC |
| I210-antisense | CCTCCTTCACCTCCAAATCMNNCGTCTGGATGCCCTCGAAG |
| T229 A230-antisense | AACACGCGGTTGCCGTAGTGGAAMNNMNNGGTCAGGATGTCGTAGGCGAAGG |
| T239-antisense | CATTATCCCGTGGGTACTTMNNGAACACGCGGTTGCCGTAG |
| K240-antisense | ACCTCTAGACCGTGGGTAMNNGGTGAACACGCGGTTGCC |

| | |
|---|---|
| **QuickChange primers for the combinations of point mutations** | |
| K202L-antisense | GATGCCCTCGAAGGGAAGCCCGCTGCCCTCGCCC |
| I207S-antisense | CTCCAAATCAATCGTCTGAGAGCCCTCGAAGGGCTTGCC |
| K202L I207S-antisense | CCTCCAAATCAATCGTCTGAGAGCCCTCGAAGGGAAGCCC |
| **pET-28a-PhoCl-6His construct** | |
| F-NcoI-PhoCl | ATATACCATGGTGATCCCTGACTACTTCAAGCAGAGCTTC |
| R-PhoCl1-linker-XhoI | GTAGCCTCGAGACCTCCACCTCCCCGTGGGTACTTGGTGAACACGC |
| **pBAD/HisB-PhoCl construct** | |
| F-XhoI-g-PhoCl | CCGAGCTCGAGTGTGATCCCTGACTAC |
| R-PhoCl*-KpnI | CATCCGCCAAAACAGCCAAGCTTGGTACCTTACCGTGGGTACTTGGTGAACAC |
| **pBAD/HisB-PhoCl-MBP construct** | |
| F-XhoI-g-PhoCl | CCGAGCTCGAGTGTGATCCCTGACTAC |
| R-PhoCl-linker-KpnI | TTCATGGTACCTCCACCTCCCCGTGGGTACTTGGTGAACAC |
| **pcDNA-NES-PhoCl-mCherry construct** | |
| quickchange-NES-HindIII-antisense | AGTAGTCAGGGATCACGCTAAGCTTGCCTCCCTGCTGCTCGTCC |
| F-HindIII-PhoCl | CTTAGAAGCTTAGCGTGATCCCTGACTACTTCAAG |
| R-PhoCl-KpnI | CCTCCGGTACCCCGTGGGTACTTGGTGAACACG |
| **pcDNA-NBid-PhoCl-CBid construct** | |
| F-NheI-NBid | AGTCAGCTAGCGCCACCATGGATTGTGAGGTCAATAACGG |
| R-BamHI-NBid overlap PCR | GGATCCCTCATCGTAGCCTTCCCAC |
| F-NBid-BamHI-PhoCl overlap PCR | GTGGGAAGGCTACGATGAGGGATCCGTGATCCCTGACTACTTCAAGC |
| R-PhoCl-KpnI-CBid overlap PCR | GATCGGTTTCCATCGGTTTGAAGGGTACCCCGTGGGTACTTGGTGAAC |
| F-KpnI-CBid overlap PCR | GGTACCCTTCAAACCGATGGAAACCGATC |
| R-CBid*-XhoI | TGACTCTCGAGTTACTTGTTCAACCTGAGCAACCACG |
| F-BamHI-PhoCl | GTCAAGGATCCGTGATCCCTGACTACTTCAAGCAGAG |
| R-PhoCl-KpnI | CCTCCGGTACCCCGTGGGTACTTGGTGAACACG |
| **pcDNA-NBid-mMaple-CBid construct** | |
| F-BamHI-mMaple | AGTAGGAATCCGTGAGCAAGGGCGAGGAGACCA |
| R-mMaple-KpnI | GCACTGGTACCCTTGTACAGCTCGTCCATGCTG |
| **pcDNA-NES-DEVD-mCardinal-NLS construct** | |
| F1-NheI-NES | CACTGGCTAGCGCCACCATGAACCTGGTGGACCTGCAGAAGAAGCTGGAGG |
| F2-NES | GACCTGCAGAAGAAGCTGGAGGAGCTGGAGCTGGACGAGCAGCAGGGATCCGCCTCCGGC |
| F3-DEVD-mCardinal | CAGGGATCCGCCTCCGGCGATGAGGTGGATGGAGCCGTGAGCAAGGGCGAGGAG |
| R1-mCardinal-NLS | CGTTTTTTTTTTGGTCCGGAGCCCTTGTACAGCTCGTCCATGCC |
| R2-NLS-NLS | GGGGTCTACTTTGCGCTTCTTTTTTGGGTCAACTTTTCGTTTTTTTTTTGGTCCGGAGCC |
| R3-NLS*-XhoI | ACTGACTCGAGTTAGGTACCTACCTTGCGTTTTTTCTTGGGGTCTACTTTGCGCTTC |

**Supplementary Table 8**. Nucleotide sequences of gene expression constructs.

| Constructs | Sequence 5'-3' |
| --- | --- |
| **LgBiT-PhoCl1-SmBiT-MBP**<br>(LgBiT is highlighted in yellow, PhoCl1 is highlighted in green, SmBiT is highlighted in cyan and MBP is highlighted in grey) | ATGGTCTTCACACTCGAAGATTTCGTTGGGGACTGGGAACAGACAGCCGCGTACAACCTGGACC<br>AAGTCCTTGAACAGGGAGGTGTGTCCAGTTTGCTGCAGAATCTCGCGGTGTCCGTAACTCCGAT<br>CCAAAGGATTGTCCGCAGCGGTGAAAATGCGCTGAAGATCGACATCCATGTCATCATCCCGTAT<br>GAAGGTCTGAGCGCGGACCAAATGGCACAGATCGAAGAAGTGTTTAAGGTGGTGTACCCTGTG<br>GATGATCATCACTTTAAGGTGATCCTGCCGTATGGCACACTGGTAATCGACGGGGTTACGCCGA<br>ACATGCTGAACTATTTCGGACGGCCGTATGAAGGCATCGCCGTGTTCGACGGCAAAAAGATCAC<br>TGTAACAGGGACCCTGTGGAACGGCAACAAAATTATCGACGAGCGCCTGATCACCCCCGACGGC<br>TCCATGCTGTTCCGAGTAACCATCAACAGCGGGAGTAGCGGAGGCGGTTCTAGCGGCGGAGAA<br>CCTGGTACCGTGATCCCTGACTACTTCAAGCAGAGCTTCCCCGAGGGCTACAGCTGGGAGCGCA<br>GCATGACCTACGAGGACGGCGGCATCTGCATCGCCACCAACGACATCACAATGGAGGGGGACA<br>GCTTCATCAACAAGATCCACTTCAAGGGCACGAACTTCCCCCCCAACGGCCCCGTGATGCAGAA<br>GAGGACCGTGGGCTGGGAGGCCAGCACCGAGAAGATGTACGAGCGCGACGGCGTGCTGAAGG<br>GCGACGTGAAGATGAAGCTGCTGCTGAAGGGCGGCGGCCACTATCGCTGCGACTACCGCACCA<br>CCTACAAGGTCAAGCAGAAGCCCGTAAAGCTGCCCGACTACCACTTCGTGGACCACCGCATCGA<br>GATCCTGAGCCACGACAAGGACTACAACAAGGTGAAGCTGTACGAGCACGCCGTGGCCCGCAA<br>CTCCACCGACAGCATGGACGAGCTGTACAAGGGTGGCAGCGGTGGCATGGTGAGCAAGGGCG<br>AGGAGACCATTACAAGCGTGATCAAGCCTGACATGAAGAACAAGCTGCGCATGGAGGGCAACG<br>TGAACGGCCACGCCTTCGTGATCGAGGGCGAGGGCAGCGGCAAGCCCTTCGAGGGCATCCAGA<br>CGATTGATTTGGAGGTGAAGGAGGGCGCCCCGCTGCCCTTCGCCTACGACATCCTGACCACCGC<br>CTTCCACTACGGCAACCGCGTGTTCACCAAGTACCCACGGTCTAGAGGTTCGTCCGGTGGCGGA<br>AGCAGTGGCGGCGTGACTGGATACAGACTTTTTGAAGAGATCCTGGAATTCGGGGGTGGAGGT<br>TCAAAAATCGAAGAAGGTAAACTGGTAATCTGGATTAACGGCGATAAAGGCTATAACGGTCTCG<br>CTGAAGTCGGTAAGAAATTCGAGAAAGATACCGGAATTAAAGTCACCGTTGAGCATCCGGATAA<br>ACTGGAAGAGAAATTCCCACAGGTTGCGGCAACTGGCGATGGCCCTGACATTATCTTCTGGGCA<br>CACGACCGCTTTGGTGGCTACGCTCAATCTGGCCTGTTGGCTGAAATCACCCCGGACAAAGCGTT<br>CCAGGACAAGCTGTATCCGTTTACCTGGGATGCCGTACGTTACAACGGCAAGCTGATTGCTTACC<br>CGATCGCTGTTGAAGCGTTATCGCTGATTTATAACAAAGATCTGCTGCCGAACCCGCCAAAAACC<br>TGGGAAGAGATCCCGGCGCTGGATAAAGAACTGAAAGCGAAAGGTAAGAGCGCGCTGATGTTC<br>AACCTGCAAGAACCGTACTTCACCTGGCCGCTGATTGCTGCTGACGGGGGTTATGCGTTCAAGT<br>ATGAAAACGGCAAGTACGACATTAAAGACGTGGGCGTGGATAACGCTGGCGCGAAAGCGGGTC<br>TGACCTTCCTGGTTGACCTGATTAAAAACAAACACATGAATGCAGACACCGATTACTCCATCGCA<br>GAAGCTGCCTTTAATAAAGGCGAAACAGCGATGACCATCAACGGCCCGTGGGCATGGTCCAACA<br>TCGACACCAGCAAAGTGAATTATGGTGTAACGGTACTGCCGACCTTCAAGGGTCAACCATCCAA<br>ACCGTTCGTTGGCGTGCTGAGCGCAGGTATTAACGCCGCCAGTCCGAACAAAGAGCTGGCAAA<br>AGAGTTCCTCGAAAACTATCTGCTGACTGATGAAGGTCTGGAAGCGGTTAATAAAGACAAACCG<br>CTGGGTGCCGTAGCGCTGAAGTCTTACGAGGAAGAGTTGGTGAAAGATCCGCGTATTGCCGCC<br>ACTATGGAAAACGCCCAGAAAGGTGAAATCATGCCGAACATCCCGCAGATGTCCGCTTTCTGGT<br>ATGCCGTGCGTACTGCGGTGATCAACGCCGCCAGCGGTCGTCAGACTGTCGATGAAGCCCTGAA<br>AGACGCGCAGACTTAA |
| **PhoCl2c**<br>(PhoCl is highlighted in green and mutations compared to PhoCl1 are highlighted in magenta) | GTGATCCCTGACTACTTCAAGCAGAGCTTCCCCGAGGGCTACAGCTGGGAGCGCAGCATGACCT<br>ACGAGGACGGCGGCATCTGCATCGCCACCAACGACATCACAATGGAGGGGGACAGCTTCATCA<br>ACAAGATCCACTTCCAGGGCACGAACTTCCCCCCCAACGGCCCCGTGATGCAGAAGAGGACCGT<br>GGGCTGGGAGGCCAGCACCGAGAAGATGTACGAGCGCGACGGCGTGCTGAAGGGCGACGTGA<br>AGATGAAGCTGCTGCTGAAGGGCGGCGGCCACTATCGCGGCGACTACCGCACCACCTACAAGG<br>TCAAGCAGAAGCCCGTAAAGCTGCCCGACTGCCACTTCGTGGACCACCGCATCGAGATCCTGAG<br>CCACGACAAGGACTACAACAAGGTGAAGCTGTACGAGCACGCCGTGGCCAAGACTTCCACCGA<br>CAGCATGGACGAGCTGTACAAGGGTGGCAGCGGTGGCATGGTGAGCAAGGGCGAGGAGACCA<br>TTACAAGCGTGATCAAGCCTGACATGAAGAACAAGCTGCGCATGGAGGGCAACGTGAACGGCC<br>ACGCCTTCGTGATCGAGGGCGAGGGCAGCGGCAAGCCCTTCGAGGGCATCCAGACGATTGATT<br>TGGAGGTGAAGGAGGGCGCCCCGCTGCCCTTCGCCTACGACATCCTGACCACCGCCTTCCACTA<br>CGGCAACCGCGTGTTCACCAAGTACCCACGG |
| **PhoCl2f**<br>(PhoCl is highlighted in green and mutations compared to PhoCl1 are highlighted in magenta) | GTGATCCCTGACTACTTCAAGCAGAGCTTCCCCGAGGGCTACAGCTGGGAGCGCAGCATGACCT<br>ACGAGGACGGCGGCATCTGCATCGCCACCAACGACATCACAATGGAGGGGGACAGCTTCATCA<br>ACAAGATCCACTTCCAGGGCACGAACTTCCCCCCCAACGGCCCCGTGATGCAGAAGAGGACCGT<br>GGGCTGGGAGGCCAGCACCGAGAAGATGTACGAGCGCGACGGCGTGCTGAAGGGCGACGTGA<br>AGATGAAGCTGCTGCTGAAGGGCGGCGGCCACTATCGCTGCGACTACCGCACCACCTACAAGGT<br>CAAGCAGAAGCCCGTAAAGCTGCCCGACTGCCACTTCGTGGACCACCGCATCGAGATCCTGAGC |

| | |
|---|---|
| | CACGACAAGGACTACAACAAGGTGAAGCTGTACGAGCACGCCGTGGCCAAGACTTCCACCGAC<br>AGCATGGACGAGCTGTACAAGGGTGGCAGCGGTGGCATGGTGAGCAAGGGCGAGGAGACCAT<br>TACAAGCGTGATCAAGCCTGACATGAAGAACAAGCTGCGCATGGAGGGCAACGTGAACGGCCA<br>CGCCTTCGTGATCGAGGGCGAGGGCAGCGGCAAGCCCTTCGAGGGCTCTCAGACGATTGATTT<br>GGAGGTGAAGGAGGGCGCCCCGCTGCCCTTCGCCTACGACATCCTGACCACCGCCTTCCACTAC<br>GGCAACCGCGTGTTCACCAAGTACCCACGG |
| **PhoCl1.5f-MV**<br>(PhoCl is highlighted in green and mutations compared to PhoCl1 are highlighted in magenta) | GTGATCCCTGACTACTTCAAGCAGAGCTTCCCCGAGGGCTACAGCTGGGAGCGCAGCATGACCT<br>ACGAGGACGGCGGCATCTGCATCGCCACCAACGACATCACAATGGAGGGGGACAGCTTCATCA<br>ACAAGATCCACTTCCAGGGCACGAACTTCCCCCCCAACGGCCCCGTGATGCAGAAGAGGACCGT<br>GGGCTGGGAGGCCAGCACCGAGAAGATGTACGAGCGCGACGGCGTGCTGAAGGGCGACGTGA<br>AGATGAAGCTGCTGCTGAAGGGCGGCGGCCACTATCGCTGCGACTACCGCACCACCTACAAGGT<br>CAAGCAGAAGCCCGTAAAGCTGCCCGACTGCCACTTCGTGGACCACCGCATCGAGATCCTGAGC<br>CACGACAAGGACTACAACAAGGTGAAGCTGTACGAGCACGCCATGGTTAAGACTTCCACCGACA<br>GCATGGACGAGCTGTACAAGGGTGGCAGCGGTGGCATGGTGAGCAAGGGCGAGGAGACCATT<br>ACAAGCGTGATCAAGCCTGACATGAAGAACAAGCTGCGCATGGAGGGCAACGTGAACGGCCAC<br>GCCTTCGTGATCGAGGGCGAGGGCAGCGGCAAGCCCTTCGAGGGCATCCAGACGATTGATTTG<br>GAGGTGAAGGAGGGCGCCCCGCTGCCCTTCGCCTACGACATCCTGACCACCGCCTTCCACTACG<br>GCAACCGCGTGTTCACCAAGTACCCACGG |
| **PhoCl1.5f-L**<br>(PhoCl is highlighted in green and mutations compared to PhoCl1 are highlighted in magenta) | GTGATCCCTGACTACTTCAAGCAGAGCTTCCCCGAGGGCTACAGCTGGGAGCGCAGCATGACCT<br>ACGAGGACGGCGGCATCTGCATCGCCACCAACGACATCACAATGGAGGGGGACAGCTTCATCA<br>ACAAGATCCACTTCCAGGGCACGAACTTCCCCCCCAACGGCCCCGTGATGCAGAAGAGGACCGT<br>GGGCTGGGAGGCCAGCACCGAGAAGATGTACGAGCGCGACGGCGTGCTGAAGGGCGACGTGA<br>AGATGAAGCTGCTGCTGAAGGGCGGCGGCCACTATCGCTGCGACTACCGCACCACCTACAAGGT<br>CAAGCAGAAGCCCGTAAAGCTGCCCGACTGCCACTTCGTGGACCACCGCATCGAGATCCTGAGC<br>CACGACAAGGACTACAACAAGGTGAAGCTGTACGAGCACGCCGTGGCCAAGACTTCCACCGAC<br>AGCATGGACGAGCTGTACAAGGGTGGCAGCGGTGGCATGGTGAGCAAGGGCGAGGAGACCAT<br>TACAAGCGTGATCAAGCCTGACATGAAGAACAAGCTGCGCATGGAGGGCAACGTGAACGGCCA<br>CGCCTTCGTGATCGAGGGCGAGGGCAGCGGGCTTCCCTTCGAGGGCATCCAGACGATTGATTTG<br>GAGGTGAAGGAGGGCGCCCCGCTGCCCTTCGCCTACGACATCCTGACCACCGCCTTCCACTACG<br>GCAACCGCGTGTTCACCAAGTACCCACGG |
| **PhoCl1.5f-MVL**<br>(PhoCl is highlighted in green and mutations compared to PhoCl1 are highlighted in magenta) | GTGATCCCTGACTACTTCAAGCAGAGCTTCCCCGAGGGCTACAGCTGGGAGCGCAGCATGACCT<br>ACGAGGACGGCGGCATCTGCATCGCCACCAACGACATCACAATGGAGGGGGACAGCTTCATCA<br>ACAAGATCCACTTCCAGGGCACGAACTTCCCCCCCAACGGCCCCGTGATGCAGAAGAGGACCGT<br>GGGCTGGGAGGCCAGCACCGAGAAGATGTACGAGCGCGACGGCGTGCTGAAGGGCGACGTGA<br>AGATGAAGCTGCTGCTGAAGGGCGGCGGCCACTATCGCTGCGACTACCGCACCACCTACAAGGT<br>CAAGCAGAAGCCCGTAAAGCTGCCCGACTGCCACTTCGTGGACCACCGCATCGAGATCCTGAGC<br>CACGACAAGGACTACAACAAGGTGAAGCTGTACGAGCACGCCATGGTTAAGACTTCCACCGACA<br>GCATGGACGAGCTGTACAAGGGTGGCAGCGGTGGCATGGTGAGCAAGGGCGAGGAGACCATT<br>ACAAGCGTGATCAAGCCTGACATGAAGAACAAGCTGCGCATGGAGGGCAACGTGAACGGCCAC<br>GCCTTCGTGATCGAGGGCGAGGGCAGCGGGCTTCCCTTCGAGGGCATCCAGACGATTGATTTG<br>GAGGTGAAGGAGGGCGCCCCGCTGCCCTTCGCCTACGACATCCTGACCACCGCCTTCCACTACG<br>GCAACCGCGTGTTCACCAAGTACCCACGG |
| **PhoCl1.5f-MVS**<br>(PhoCl is highlighted in green and mutations compared to PhoCl1 are highlighted in magenta) | GTGATCCCTGACTACTTCAAGCAGAGCTTCCCCGAGGGCTACAGCTGGGAGCGCAGCATGACCT<br>ACGAGGACGGCGGCATCTGCATCGCCACCAACGACATCACAATGGAGGGGGACAGCTTCATCA<br>ACAAGATCCACTTCCAGGGCACGAACTTCCCCCCCAACGGCCCCGTGATGCAGAAGAGGACCGT<br>GGGCTGGGAGGCCAGCACCGAGAAGATGTACGAGCGCGACGGCGTGCTGAAGGGCGACGTGA<br>AGATGAAGCTGCTGCTGAAGGGCGGCGGCCACTATCGCTGCGACTACCGCACCACCTACAAGGT<br>CAAGCAGAAGCCCGTAAAGCTGCCCGACTGCCACTTCGTGGACCACCGCATCGAGATCCTGAGC<br>CACGACAAGGACTACAACAAGGTGAAGCTGTACGAGCACGCCATGGTTAAGACTTCCACCGACA<br>GCATGGACGAGCTGTACAAGGGTGGCAGCGGTGGCATGGTGAGCAAGGGCGAGGAGACCATT<br>ACAAGCGTGATCAAGCCTGACATGAAGAACAAGCTGCGCATGGAGGGCAACGTGAACGGCCAC<br>GCCTTCGTGATCGAGGGCGAGGGCAGCGGCAAGCCCTTCGAGGGCTCTCAGACGATTGATTTG<br>GAGGTGAAGGAGGGCGCCCCGCTGCCCTTCGCCTACGACATCCTGACCACCGCCTTCCACTACG<br>GCAACCGCGTGTTCACCAAGTACCCACGG |
| **PhoCl1.5f-LS**<br>(PhoCl is highlighted in green and mutations compared to PhoCl1 are highlighted in magenta) | GTGATCCCTGACTACTTCAAGCAGAGCTTCCCCGAGGGCTACAGCTGGGAGCGCAGCATGACCT<br>ACGAGGACGGCGGCATCTGCATCGCCACCAACGACATCACAATGGAGGGGGACAGCTTCATCA<br>ACAAGATCCACTTCCAGGGCACGAACTTCCCCCCCAACGGCCCCGTGATGCAGAAGAGGACCGT<br>GGGCTGGGAGGCCAGCACCGAGAAGATGTACGAGCGCGACGGCGTGCTGAAGGGCGACGTGA<br>AGATGAAGCTGCTGCTGAAGGGCGGCGGCCACTATCGCTGCGACTACCGCACCACCTACAAGGT<br>CAAGCAGAAGCCCGTAAAGCTGCCCGACTGCCACTTCGTGGACCACCGCATCGAGATCCTGAGC<br>CACGACAAGGACTACAACAAGGTGAAGCTGTACGAGCACGCCGTGGCCAAGACTTCCACCGAC<br>AGCATGGACGAGCTGTACAAGGGTGGCAGCGGTGGCATGGTGAGCAAGGGCGAGGAGACCAT |

| | |
|---|---|
| | TACAAGCGTGATCAAGCCTGACATGAAGAACAAGCTGCGCATGGAGGGCAACGTGAACGGCCA<br>CGCCTTCGTGATCGAGGGCGAGGGCAGCGGGCTTCCCTTCGAGGGCTCTCAGACGATTGATTTG<br>GAGGTGAAGGAGGGCGCCCCGCTGCCCTTCGCCTACGACATCCTGACCACCGCCTTCCACTACG<br>GCAACCGCGTGTTCACCAAGTACCCACGG |
| **PhoCl1.5f-MVLS**<br>(PhoCl is highlighted in green and mutations compared to PhoCl1 are highlighted in magenta) | GTGATCCCTGACTACTTCAAGCAGAGCTTCCCCGAGGGCTACAGCTGGGAGCGCAGCATGACCT<br>ACGAGGACGGCGGCATCTGCATCGCCACCAACGACATCACAATGGAGGGGGACAGCTTCATCA<br>ACAAGATCCACTTCCAGGGCACGAACTTCCCCCCCAACGGCCCCGTGATGCAGAAGAGGACCGT<br>GGGCTGGGAGGCCAGCACCGAGAAGATGTACGAGCGCGACGGCGTGCTGAAGGGCGACGTGA<br>AGATGAAGCTGCTGCTGAAGGGCGGCGGCCACTATCGCTGCGACTACCGCACCACCTACAAGGT<br>CAAGCAGAAGCCCGTAAAGCTGCCCGACTGCCACTTCGTGGACCACCGCATCGAGATCCTGAGC<br>CACGACAAGGACTACAACAAGGTGAAGCTGTACGAGCACGCCATGGTTAAGACTTCCACCGACA<br>GCATGGACGAGCTGTACAAGGGTGGCAGCGGTGGCATGGTGAGCAAGGGCGAGGAGACCATT<br>ACAAGCGTGATCAAGCCTGACATGAAGAACAAGCTGCGCATGGAGGGCAACGTGAACGGCCAC<br>GCCTTCGTGATCGAGGGCGAGGGCAGCGGGCTTCCCTTCGAGGGCTCTCAGACGATTGATTTGG<br>AGGTGAAGGAGGGCGCCCCGCTGCCCTTCGCCTACGACATCCTGACCACCGCCTTCCACTACGG<br>CAACCGCGTGTTCACCAAGTACCCACGG |
| **PhoCl1-6His**<br>(PhoCl is highlighted in green and His tag is highlighted in yellow) | ATGGTGATCCCTGACTACTTCAAGCAGAGCTTCCCCGAGGGCTACAGCTGGGAGCGCAGCATGA<br>CCTACGAGGACGGCGGCATCTGCATCGCCACCAACGACATCACAATGGAGGGGGACAGCTTCAT<br>CAACAAGATCCACTTCAAGGGCACGAACTTCCCCCCCAACGGCCCCGTGATGCAGAAGAGGACC<br>GTGGGCTGGGAGGCCAGCACCGAGAAGATGTACGAGCGCGACGGCGTGCTGAAGGGCGACGT<br>GAAGATGAAGCTGCTGCTGAAGGGCGGCGGCCACTATCGCTGCGACTACCGCACCACCTACAA<br>GGTCAAGCAGAAGCCCGTAAAGCTGCCCGACTACCACTTCGTGGACCACCGCATCGAGATCCTG<br>AGCCACGACAAGGACTACAACAAGGTGAAGCTGTACGAGCACGCCGTGGCCCGCAACTCCACC<br>GACAGCATGGACGAGCTGTACAAGGGTGGCAGCGGTGGCATGGTGAGCAAGGGCGAGGAGAC<br>CATTACAAGCGTGATCAAGCCTGACATGAAGAACAAGCTGCGCATGGAGGGCAACGTGAACGG<br>CCACGCCTTCGTGATCGAGGGCGAGGGCAGCGGCAAGCCCTTCGAGGGCATCCAGACGATTGA<br>TTTGGAGGTGAAGGAGGGCGCCCCGCTGCCCTTCGCCTACGACATCCTGACCACCGCCTTCCACT<br>ACGGCAACCGCGTGTTCACCAAGTACCCACGGGGAGGTGGAGGTCTCGAGCACCACCACCACC<br>ACCACTGA |
| **NBid-PhoCl1-CBid**<br>(NBid is highlighted in yellow, PhoCl is highlighted in green and CBid is highlighted in blue) | ATGGATTGTGAGGTCAATAACGGTTCATCTCTCCGAGACGAATGCATAACGAACTTGCTCGTTTT<br>CGGCTTTCTGCAATCCTGCAGCGATAATTCTTTCAGAAGAGAACTTGACGCTCTTGGACATGAAC<br>TCCCAGTACTCGCTCCACAGTGGGAAGGCTACGATGAGGGATCCGTGATCCCTGACTACTTCAA<br>GCAGAGCTTCCCCGAGGGCTACAGCTGGGAGCGCAGCATGACCTACGAGGACGGCGGCATCTG<br>CATCGCCACCAACGACATCACAATGGAGGGGGACAGCTTCATCAACAAGATCCACTTCAAGGGC<br>ACGAACTTCCCCCCCAACGGCCCCGTGATGCAGAAGAGGACCGTGGGCTGGGAGGCCAGCACC<br>GAGAAGATGTACGAGCGCGACGGCGTGCTGAAGGGCGACGTGAAGATGAAGCTGCTGCTGAA<br>GGGCGGCGGCCACTATCGCTGCGACTACCGCACCACCTACAAGGTCAAGCAGAAGCCCGTAAA<br>GCTGCCCGACTACCACTTCGTGGACCACCGCATCGAGATCCTGAGCCACGACAAGGACTACAAC<br>AAGGTGAAGCTGTACGAGCACGCCGTGGCCCGCAACTCCACCGACAGCATGGACGAGCTGTAC<br>AAGGGTGGCAGCGGTGGCATGGTGAGCAAGGGCGAGGAGACCATTACAAGCGTGATCAAGCC<br>TGACATGAAGAACAAGCTGCGCATGGAGGGCAACGTGAACGGCCACGCCTTCGTGATCGAGGG<br>CGAGGGCAGCGGCAAGCCCTTCGAGGGCATCCAGACGATTGATTTGGAGGTGAAGGAGGGCG<br>CCCCGCTGCCCTTCGCCTACGACATCCTGACCACCGCCTTCCACTACGGCAACCGCGTGTTCACCA<br>AGTACCCACGGGGTACCCTTCAAACCGATGGAAACCGATCATCTCATTCAAGGTTGGGGCGAAT<br>TGAAGCAGATAGTGAGAGTCAGGAGGACATAATACGCAATATAGCTCGACACCTTGCACAGGTC<br>GGTGATAGCATGGACCGCTCTATCCCCCCAGGTTTGGTTAACGGATTGGCCCTGCAACTGCGAA<br>ACACTTCAAGGAGTGAAGAAGATAGAAATCGAGACCTGGCGACCGCGCTTGAACAGCTTCTTCA<br>AGCCTATCCCAGAGATATGGAGAAGGAAAAAACAATGCTCGTGCTCGCACTGCTGCTGGCAAAG<br>AAAGTAGCCTCTAACACACCATCCCTCTTGAGAGACGTCTTCCATACCACGGTAAATTTCATAAAC<br>CAGAACCTTAGGACGTATGTGCGGAGTCTTGCTCGCAATGGTATGACTGAGTTTCTTCAACCCGT<br>GCCCCTCAATGCGTGGTTGCTCAGGTTGAACAAGTAA |
| **NBid-mMaple-CBid**<br>(NBid is highlighted in yellow, mMaple is highlighted in green and CBid is highlighted in blue) | ATGGATTGTGAGGTCAATAACGGTTCATCTCTCCGAGACGAATGCATAACGAACTTGCTCGTTTT<br>CGGCTTTCTGCAATCCTGCAGCGATAATTCTTTCAGAAGAGAACTTGACGCTCTTGGACATGAAC<br>TCCCAGTACTCGCTCCACAGTGGGAAGGCTACGATGAGGGATCC<br>GTGAGCAAGGGCGAGGAGACCATTATGAGCGTGATCAAGCCTGACATGAAGATCAAGCTGCGC<br>ATGGAGGGCAACGTGAACGGCCACGCCTTCGTGATCGAGGGCGAGGGCAGCGGCAAGCCCTTC<br>GAGGGCATCCAGACGATTGATTTGGAGGTGAAGGAGGGCGCCCCGCTGCCCTTCGCCTACGAC<br>ATCCTGACCACCGCCTTCCACTACGGCAACCGCGTGTTCACCAAGTACCCCGAGGACATCCCTGA<br>CTACTTCAAGCAGAGCTTCCCCGAGGGCTACAGCTGGGAGCGCAGCATGACCTACGAGGACGG<br>CGGCATCTGCATCGCCACCAACGACATCACAATGGAGGAGGACAGCTTCATCAACAAGATCCAC<br>TTCAAGGGCACGAACTTCCCCCCCAACGGCCCCGTGATGCAGAAGAGGACCGTGGGCTGGGAG<br>GTCAGCACCGAGAAGATGTACGTGCGCGACGGCGTGCTGAAGGGCGACGTGAAGATGAAGCT |

| | |
|---|---|
| | GCTGCTGAAGGGCGGCAGCCACTATCGCTGCGACTTCCGCACCACCTACAAGGTCAAGCAGAAG<br>GCCGTAAAGCTGCCCGACTACCACTTCGTGGACCACCGCATCGAGATCCTGAGCCACGACAAGG<br>ACTACAACAAGGTGAAGCTGTACGAGCACGCCGTGGCCCGCAACTCCACCGACAGCATGGACG<br>AGCTGTACAAGGGTACCCTTCAAACCGATGGAAACCGATCATCTCATTCAAGGTTGGGGCGAAT<br>TGAAGCAGATAGTGAGAGTCAGGAGGACATAATACGCAATATAGCTCGACACCTTGCACAGGTC<br>GGTGATAGCATGGACCGCTCTATCCCCCCAGGTTTGGTTAACGGATTGGCCCTGCAACTGCGAA<br>ACACTTCAAGGAGTGAAGAAGATAGAAATCGAGACCTGGCGACCGCGCTTGAACAGCTTCTTCA<br>AGCCTATCCCAGAGATATGGAGAAGGAAAAAACAATGCTCGTGCTCGCACTGCTGCTGGCAAAG<br>AAAGTAGCCTCTAACACACCATCCCTCTTGAGAGACGTCTTCCATACCACGGTAAATTTCATAAAC<br>CAGAACCTTAGGACGTATGTGCGGAGTCTTGCTCGCAATGGTATGACTGAGTTTCTTCAACCCGT<br>GCCCCTCAATGCGTGGTTGCTCAGGTTGAACAAGTAA |
| **NBid-PhoCl2c-CBid**<br>(NBid is highlighted in yellow, PhoCl2c is highlighted in green and CBid is highlighted in blue) | ATGGATTGTGAGGTCAATAACGGTTCATCTCTCCGAGACGAATGCATAACGAACTTGCTCGTTTT<br>CGGCTTTCTGCAATCCTGCAGCGATAATTCTTTCAGAAGAGAACTTGACGCTCTTGGACATGAAC<br>TCCCAGTACTCGCTCCACAGTGGGAAGGCTACGATGAGGGATCCGTGATCCCTGACTACTTCAA<br>GCAGAGCTTCCCCGAGGGCTACAGCTGGGAGCGCAGCATGACCTACGAGGACGGCGGCATCTG<br>CATCGCCACCAACGACATCACAATGGAGGGGGACAGCTTCATCAACAAGATCCACTTCCAGGGC<br>ACGAACTTCCCCCCCAACGGCCCCGTGATGCAGAAGAGGACCGTGGGCTGGGAGGCCAGCACC<br>GAGAAGATGTACGAGCGCGACGGCGTGCTGAAGGGCGACGTGAAGATGAAGCTGCTGCTGAA<br>GGGCGGCGGCCACTATCGCGGCGACTACCGCACCACCTACAAGGTCAAGCAGAAGCCCGTAAA<br>GCTGCCCGACTGCCACTTCGTGGACCACCGCATCGAGATCCTGAGCCACGACAAGGACTACAAC<br>AAGGTGAAGCTGTACGAGCACGCCGTGGCCAAGACTTCCACCGACAGCATGGACGAGCTGTAC<br>AAGGGTGGCAGCGGTGGCATGGTGAGCAAGGGCGAGGAGACCATTACAAGCGTGATCAAGCC<br>TGACATGAAGAACAAGCTGCGCATGGAGGGCAACGTGAACGGCCACGCCTTCGTGATCGAGGG<br>CGAGGGCAGCGGCAAGCCCTTCGAGGGCATCCAGACGATTGATTTGGAGGTGAAGGAGGGCG<br>CCCCGCTGCCCTTCGCCTACGACATCCTGACCACCGCCTTCCACTACGGCAACCGCGTGTTCACCA<br>AGTACCCACGGGGTACCCTTCAAACCGATGGAAACCGATCATCTCATTCAAGGTTGGGGCGAAT<br>TGAAGCAGATAGTGAGAGTCAGGAGGACATAATACGCAATATAGCTCGACACCTTGCACAGGTC<br>GGTGATAGCATGGACCGCTCTATCCCCCCAGGTTTGGTTAACGGATTGGCCCTGCAACTGCGAA<br>ACACTTCAAGGAGTGAAGAAGATAGAAATCGAGACCTGGCGACCGCGCTTGAACAGCTTCTTCA<br>AGCCTATCCCAGAGATATGGAGAAGGAAAAAACAATGCTCGTGCTCGCACTGCTGCTGGCAAAG<br>AAAGTAGCCTCTAACACACCATCCCTCTTGAGAGACGTCTTCCATACCACGGTAAATTTCATAAAC<br>CAGAACCTTAGGACGTATGTGCGGAGTCTTGCTCGCAATGGTATGACTGAGTTTCTTCAACCCGT<br>GCCCCTCAATGCGTGGTTGCTCAGGTTGAACAAGTAA |
| **NES-DEVD-mCardinal-NLS**<br>(NES is highlighted in yellow, DEVD is highlighted in green, mCardinal is highlighted in magenta and 3x NLS is highlighted in cyan) | ATGAACCTGGTGGACCTGCAGAAGAAGCTGGAGGAGCTGGAGCTGGACGAGCAGCAGGGATC<br>CGCCTCCGGCGATGAGGTGGATGGAGCCGTGAGCAAGGGCGAGGAGCTGATCAAGGAGAACA<br>TGCACATGAAGCTGTACATGGAAGGCACCGTGAACAACCACCACTTCAAGTGCACCACCGAAGG<br>GGAGGGCAAGCCCTACGAGGGCACCCAGACCCAGAGGATTAAGGTGGTGGAGGGAGGCCCCC<br>TGCCGTTCGCATTCGACATCCTGGCCACCTGCTTTATGTACGGGAGCAAGACCTTCATCAACCAC<br>ACCCAGGGCATCCCCGATTTCTTTAAGCAGTCCTTCCCTGAGGGCTTCACATGGGAGAGAGTCAC<br>CACATACGAAGACGGGGGCGTGCTTACCGTTACCCAGGACACCAGCCTCCAGGACGGCTGCTTG<br>ATCTACAACGTCAAGCTCAGAGGGGTGAACTTCCCATCCAACGGCCCTGTGATGCAGAAGAAAA<br>CACTCGGCTGGGAGGCCACCACCGAGACCCTGTACCCCGCTGACGGCGGCCTGGAAGGCAGAT<br>GCGACATGGCCCTGAAGCTCGTGGGCGGGGGCCACCTGCACTGCAACCTGAAGACCACATACA<br>GATCCAAGAAACCCGCTAAGAACCTCAAGATGCCCGGCGTCTACTTTGTGGACCGCAGACTGGA<br>AAGAATCAAGGAGGCCGACAATGAGACCTACGTCGAGCAGCACGAGGTGGCTGTGGCCAGATA<br>CTGCGACCTCCCTAGCAAACTGGGGCACAAACTTAATGGCATGGACGAGCTGTACAAGGGCTCC<br>GGACCAAAAAAAAAAACGAAAAGTTGACCCAAAAAAGAAGCGCAAAGTAGACCCCAAGAAAAAA<br>CGCAAGGTAGGTACCTAA |

### References:

1. Wallace, A. C., Laskowski, R. A. & Thornton, J. M. LIGPLOT: a program to generate schematic diagrams of protein-ligand interactions. *Protein Eng.* **8**, 127–134 (1995).

2. Shaner, N. C., Steinbach, P. A. & Tsien, R. Y. A guide to choosing fluorescent proteins. *Nat. Methods* **2**, 905–909 (2005).

3. Heim, R., Cubitt, A. B. & Tsien, R. Y. Improved green fluorescence. *Nature* **373**, 663–664 (1995).
